# The H3^K36M^ oncohistone inhibits NSD2 to activate a SETD2-dependent antiviral-like immune response in KRAS-driven lung cancer

**DOI:** 10.1101/2025.05.25.655971

**Authors:** Amy C. Gladstein, Carson D. Poltorack, Akino Mercy Charles Solomon, Sharan Venkatesh, Keren M. Adler, Maggie R. Robertson, Stephanie Stransky, Valerie M. Irizarry-Negron, Dain A. Ruiz, Nelson F. Freeburg, Simone Sidoli, Irfan A. Asangani, Sydney M. Shaffer, David M. Feldser

## Abstract

Mutations in histone 3 at or near lysine 36 (H3K36) have dominantly acting oncogenic effects in multiple tumor types by limiting H3K36-directed methyltransferases. Paradoxically, we find that expression of the *H3^K36M^* oncohistone unexpectedly inhibits tumor formation in KRAS-driven lung adenocarcinoma by inducing a potent immune-mediated tumor clearance. Mechanistically, oncohistone expression derepresses endogenous retroviral element transcription, results in the accumulation of double-stranded RNA (dsRNA), and activates an innate antiviral-like immune response that eradicates tumor growth. Surprisingly, while inactivation of the H3K36 di-methyltransferase NSD2 replicated all effects of oncohistone expression, inactivation of the H3K36 tri-methyltransferase SETD2 abolished element derepression and all associated downstream anti-cancer effects that are induced by oncohistone expression. These observations restructure our understanding of the roles of H3K36 methylation, the consequences of its deregulation in cancer, and shape our expectations for therapeutic interventions targeting H3K36 methyltransferases.

## INTRODUCTION

Genes encoding histone proteins (*H3.1* and *H3.3*) are mutated at or near lysine 36 in nearly all cases of chondroblastoma, frequently in undifferentiated sarcomas, and in head and neck squamous cell carcinomas [1]. These mutations have been dubbed ‘oncohistones’ because they are causally linked to cancer formation, expressed in the context of other wildtype *H3* alleles, and act dominantly to inhibit histone H3 lysine 36 (H3K36) methylation both in cis at chromatin sites where they are incorporated and more broadly in trans due to widespread inhibition of H3K36 methyltransferases [2, 3]. H3K36 methylation is a critical post-translational histone modification that regulates genomic stability, transcriptional fidelity, and cell identity [4, 5]. In addition to oncohistone mutations, mutations in H3K36 methyltransferases are a common feature of diverse cancer types [6]. H3K36 can be mono-, di-, or tri-methylated, with each methylation state having distinct chromatin localization and functional effects [4]. Multiple enzymes, such as NSD1 and NSD2, catalyze the mono– and di-methylation states of H3K36, while SETD2 is the only enzyme known to catalyze all three methylation states [4].

Inactivation of *SETD2* is prevalent in a wide range of malignancies, including clear cell renal cell carcinoma, gliomas, and KRAS-driven lung adenocarcinoma [6]. *SETD2* is a potent tumor suppressor in KRAS-driven lung adenocarcinoma, with its inactivation promoting rapid acceleration of tumor cell proliferation [7–10]. SETD2 binds to RNA Polymerase II and co-transcriptionally promotes H3K36 trimethylation (H3K36me3) [11, 12]. While H3K36me3 marks actively transcribed genes, it is thought to facilitate resetting of chromatin to a repressed state after transcription [13, 14]. H3K36me3 also regulates diverse cellular processes, including DNA repair, alternative splicing, and RNA processing [14–18]. More recently, SETD2 has also been found to methylate non-histone substrates, playing critical roles in cytoskeletal remodeling (α-tubulin and actin) [19, 20], interferon signaling (STAT1) [21], and transcriptional repression (EZH2) [22]. While SETD2 is frequently inactivated in diverse malignancies, the H3K36 di-methyltransferases NSD1 and NSD2 are frequently over-expressed in cancer [23]. Notably, NSD2 acts as a driver of tumor progression in KRAS-driven lung adenocarcinoma [24]. H3K36 dimethylation (H3K36me2) is specifically enriched in intergenic regions and has known roles in DNA repair and DNA methylation, but it remains relatively understudied [25, 26]. Importantly, both H3K36me2 and H3K36me3 impair the action of PRC2, the enzyme that catalyzes H3K27me3 and facilitates transcriptional silencing at chromatin regions where it is enriched [27].

Although both *SETD2* inactivation and oncohistone expression disrupt H3K36 methylation, *SETD2* is frequently inactivated in lung adenocarcinoma, whereas *H3^K36M^*, or other oncohistone mutations, have not been observed in this cancer type [1]. Therefore, we sought to understand the effects of expressing the *H3^K36M^* oncohistone in KRAS*-*driven lung adenocarcinoma. Using KRAS-driven mouse models of lung adenocarcinoma, we found that *H3^K36M^* expression is detrimental to tumor growth, but that this effect is entirely dependent on SETD2 function. Mechanistically, we uncovered that oncohistone expression results in the derepression of endogenous retroviral elements, promotes the subsequent accumulation of dsRNA, which culminates in a RIG-I/MDA5-mediated anti-tumor immune response. Surprisingly, inactivation of *Setd2* abrogates retroviral element expression and accumulation of dsRNA, which in turn cancels all of the downstream antiviral-like, pro-inflammatory consequences of oncohistone expression. Moreover, when combined, inactivation of *Setd2* cooperates with oncohistone expression to further exacerbate tumor growth, revealing the oncogenic potential of oncohistone expression in this context. The opposing effects of oncohistone expression and *Setd2* inactivation are explained by the propensity of oncohistones to inhibit the H3K36 di-methyltransferase NSD2, inactivation of which phenocopies all aspects of oncohistone expression. Collectively, our findings define a molecular mechanism by which distinct H3K36 methyltransferases function in cancer and implicate NSD2 as an anti-cancer target in KRAS-driven lung adenocarcinoma.

## RESULTS

### H3^K36M^ limits KRAS-driven tumor growth in a Setd2-dependent manner

To investigate the effects of *H3^K36M^* expression in KRAS-driven lung adenocarcinoma, we expressed *H3^K36M^* in a *Kras^G12D^*-driven mouse model. *Kras^LSL-G12D/+^*; *(Rosa26^LSL-YFP^)* (*K(Y)*) and *Kras^LSL-G12D/+^*; (*Rosa26^LSL-YFP^)*; *Tg:CAGsLSL-H3^K36M^*(*K(Y)H3^K36M^*) mice were transduced with a lentiviral vector that expresses Cre, Cas9, and a *Setd2*-targeting or inert non-targeting sgRNA. This experimental scheme established four cohorts: *K-Ctrl*, *K-Setd2^KO^*, *KH3^K36M^-Ctrl*, and *KH3^K36M^-Setd2^KO^* (**Figure 1A**). As expected, *Setd2* inactivation (*K-Setd2^KO^*) significantly increased tumor size compared to *K-Ctrl* tumors (**Figures 1B, 1C, and S1A**). However, in stark contrast, expression of the oncohistone substantially reduced the number of tumors observed and the size of each tumor. Most surprisingly, when *H3^K36M^* expression was combined with *Setd2* inactivation (*KH3^K36M^-Setd2^KO^*), the number of tumors observed was restored, and the tumors were even larger than when *Setd2* was inactivated alone.

**Figure 1:**
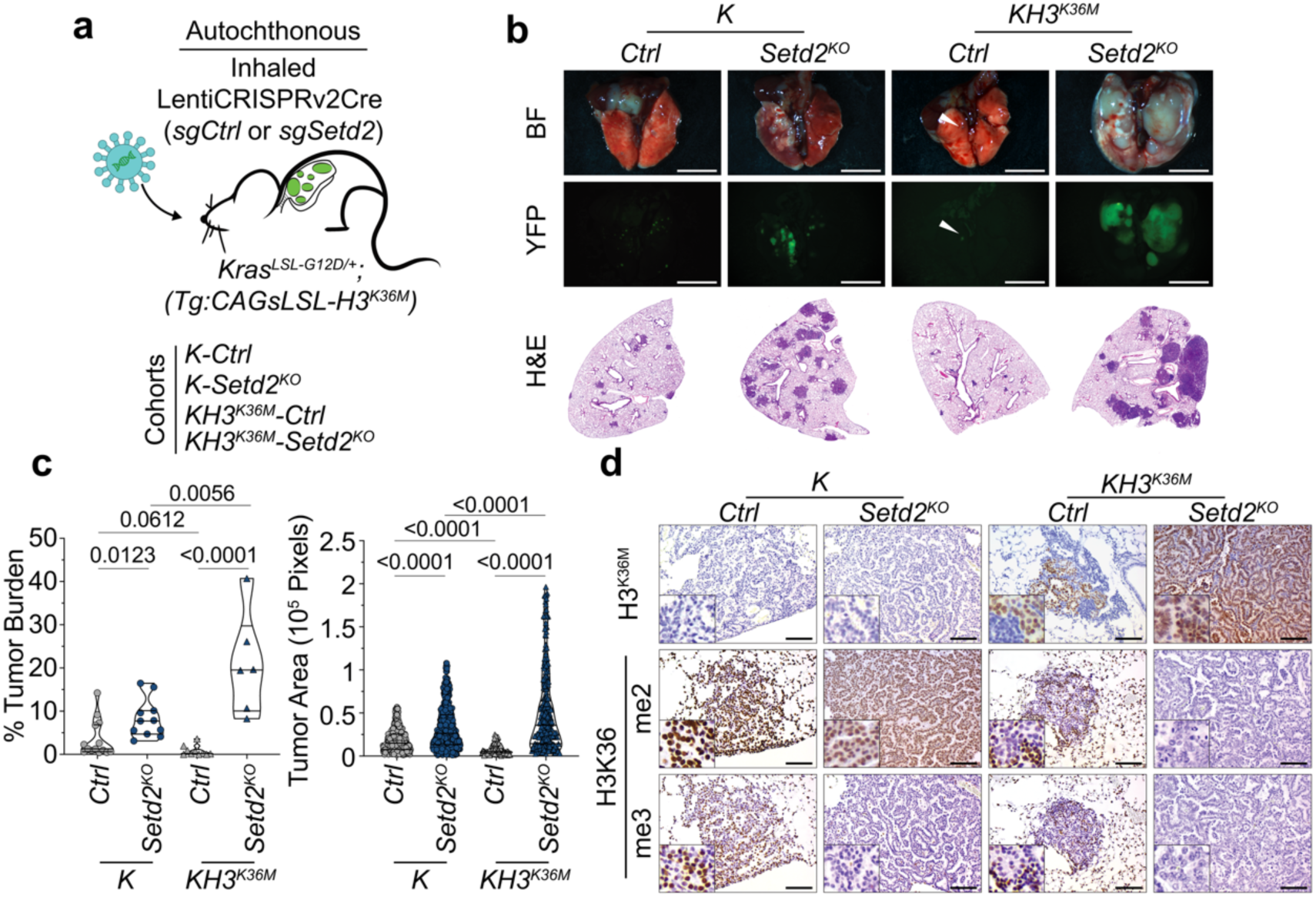
H3^K36M^ limits Kras-driven tumor growth in a Setd2-dependent manner. a. Experimental scheme of autochthonous tumor initiation. b. Brightfield and fluorescent micrographs of lungs from *K-Ctrl*, *K-Setd2^KO^*, *KH3^K36M^-Ctrl*, and *KH3^K36M^-Setd2^KO^* mice. Scale bars are 4.4 mm. Below are representative H&E images of tumor-bearing lobes from *K-Ctrl*, *K-Setd2^KO^*, *KH3^K36M^-Ctrl*, and *KH3^K36M^-Setd2^KO^* mice. c. Quantification of tumor burden and individual tumor area in *K-Ctrl*, *K-Setd2^KO^*, *KH3^K36M^-Ctrl*, and *KH3^K36M^-Setd2^KO^* mice. Significance was determined by unpaired Student’s t-test. Error bars represent mean±SEM. d. IHC for H3^K36M^, H3K36me2, and H3K36me3 in *K-Ctrl*, *K-Setd2^KO^*, *KH3^K36M^-Ctrl*, and *KH3^K36M^-Setd2^KO^* mice. Scale bars are 119 μm; insets are magnified x3.

*H3^K36M^* expression was robust in both *KH3^K36M^-Ctrl and KH3^K36M^-Setd2^KO^* tumors (**Figure 1D**). As expected, H3K36me3 was reduced in *K-Setd2^KO^*, *KH3^K36M^-Ctrl*, and *KH3^K36M^-Setd2^KO^* tumors, while H3K36me2 was reduced in *KH3^K36M^-Ctrl* and *KH3^K36M^-Setd2^KO^* tumors (**Figure 1D**). Interestingly, *KH3^K36M^-Ctrl* tumors had significantly more instances of cell death whereas *KH3^K36M^-Setd2^KO^* tumors had significantly more proliferative tumor cells compared to other tumor genotypes (**Figures S1B-E**). All tumors had typical markers of cell identity for lung adenocarcinoma cells (**Figure S1F)**. Thus, despite its impact on inhibiting H3K36 methylation, *H3^K36M^*oncohistone expression is not oncogenic and does not phenocopy the effects of *Setd2* inactivation in KRAS-driven lung cancer, but instead suppresses tumor formation and growth in a *Setd2*-dependent manner.

### H3^K36M^ promotes a Setd2-dependent tumor-cell-killing immune response

Pathology review of *KH3^K36M^-Ctrl* tumors showed a significant influx of immune cells in each tumor mass, which was absent in *KH3^K36M^-Setd2^KO^* tumors (**Figure S2A**). To better understand the frequency, repertoire, and biology of tumor-immune cell interactions within *KH3^K36M^-Ctrl* tumors, we performed spatial transcriptomics using the Xenium 5000-plex pan-tissue and signaling pathway panel on lung lobes from *K-Ctrl, K-Setd2^KO^, KH3^K36M^-Ctrl, and KH3^K36M^-Setd2^KO^* mice [28]. After quality control, we analyzed nearly 1.3 million single cells from 159 tumors and uncovered profound differences in the cellular composition, cellular states, and tissue organization of these tumors (**Figures 2A and S3A-C**). Though unremarkable from typical pathology analysis, with insight from spatial gene expression analysis, we detected a low-level immune reaction present in *K-Ctrl* tumors that was absent in *K-Setd2^KO^* tumors. As expected though, this immune reaction was significantly enhanced in *KH3^K36M^-Ctrl* tumors and absent in *KH3^K36M^-Setd2^KO^*tumors. Interestingly, the reaction was dominated by T cells, B cells, and plasma cells (**Figure 2B**). Orthogonally, we sought to confirm these observations by staining tumors with a panel of immune cell markers. Indeed, in *KH3^K36M^-Ctrl* tumors, there was a dramatic increase in CD45-positive, CD3-positive, B220-positive, and NKp46-positive cells relative to *K-Ctrl* tumors, and this immune infiltration was ablated in *K-Setd2^KO^ and KH3^K36M^-Setd2^KO^* tumors (**Figures S2B and S2C**). Additionally, the abundance of other innate immune cell types, such as macrophages and neutrophils, was similar across all tumor genotypes (**Figures S2B and S2C**). These analyses demonstrate that *Setd2* inactivation eliminates both the low-level immune reaction present in KRAS-driven tumors and the massive immune reaction that is associated with oncohistone expression. Thus, a major feature of SETD2-mediated tumor suppression is to limit immune-mediated tumor cell killing.

**Figure 2:**
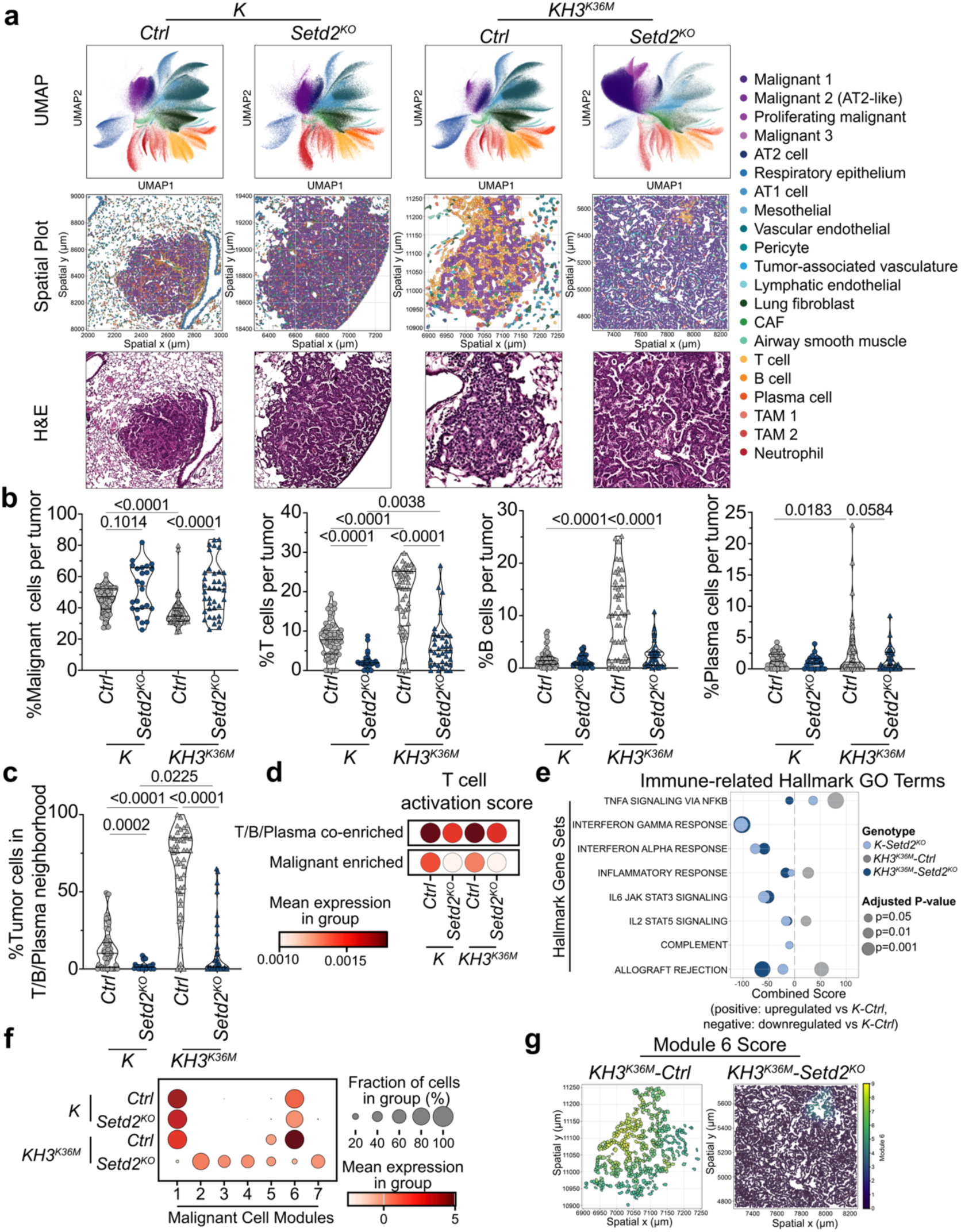
H3^K36M^ promotes a Setd2-dependent tumor-cell-killing immune response. a. UMAP embeddings of cells from *K-Ctrl*, *K-Setd2^KO^*, *KH3^K36M^-Ctrl*, and *KH3^K36M^-Setd2^KO^* lung lobes, paired with representative single-tumor spatial plots and H&E images. b. Percentage of cells within individual tumors identified as malignant, T, B, or plasma cells in cell typing. Significance was determined by unpaired Student’s t-test. Error bars represent mean±SEM. c. Percentage of cells within individual *K-Ctrl*, *K-Setd2^KO^*, *KH3^K36M^-Ctrl*, and *KH3^K36M^-Setd2^KO^* tumors belonging to the T/B/Plasma-enriched cellular neighborhood. Significance was determined by unpaired Student’s t-test. Error bars represent mean±SEM. d. T cell activation scores for T cells present in the T/B/Plasma-enriched or Malignant-enriched cellular neighborhoods in *K-Ctrl*, *K-Setd2^KO^*, *KH3^K36M^-Ctrl*, and *KH3^K36M^-Setd2^KO^* lung lobes. e. Gene Ontology enrichment scores of differentially upregulated and downregulated genes in *K-Setd2^KO^*, *KH3^K36M^-Ctrl*, and *KH3^K36M^-Setd2^KO^* tumors compared to *K-Ctrl* tumors. Enrichment for a GO Term in differentially upregulated genes is depicted on the positive x-axis while enrichment in downregulated genes is depicted on the negative x-axis. Non-significant enrichments are not plotted. f. Hotspot malignant cell module scores in *K-Ctrl*, *K-Setd2^KO^*, *KH3^K36M^-Ctrl*, and *KH3^K36M^-Setd2^KO^* lung lobes. g. Spatial plots of malignant cells colored by their expression of Hotspot Module 6 in representative *KH3^K36M^-Ctrl* and *KH3^K36M^-Setd2^KO^* tumors.

To identify phenotypic cell-to-cell interactions between malignant and immune cell types across tumor genotypes, we performed cellular neighborhood analysis [29] (**Figures S3D-F**). A cellular neighborhood comprised of T, B, and plasma cells was present in *K-Ctrl* tumors and was significantly enhanced in *KH3^K36M^*-*Ctrl* tumors (**Figure 2C**). In both contexts, *Setd2* inactivation severely suppressed the presence of the T/B/Plasma cell neighborhood. Indeed, T cells in both *K-Ctrl* and *KH3^K36M^-Ctrl* tumors had higher expression of a T cell activation gene signature than corresponding *Setd2^KO^* tumors, suggesting a more robust T cell response (**Figure 2D**). Next, we assessed malignant cell gene expression programs across tumor genotypes. We noted that Gene Ontology (GO) terms relating to inflammatory gene expression programs were significantly suppressed by *Setd2* inactivation in both *K-Setd2^KO^* and *KH3^K36M^-Setd2^KO^* tumors. Additionally, a subset of these gene expression programs was significantly enriched in oncohistone-expressing *KH3^K36M^-Ctrl* tumors relative to *K-Ctrl* tumors (**Figure 2E**). Consistent with their robust tumor growth properties, terms relating to aggressive tumor growth were enriched in *KH3^K36M^-Setd2^KO^* tumors (**Figure S4A**). Finally, to understand whether cancer cell gene expression covaries spatially with immune infiltrates, we performed Hotspot analysis on malignant cells [30]. Of the modules identified, Module 6 was noted as upregulated in *K-Ctrl* tumors, further enhanced in *KH3^K36M^-Ctrl* tumors, but reduced in both contexts by *Setd2* inactivation (**Figures 2F and 2G**). The genes in this module were predominantly immune-related and included members of the complement pathway (*C3*, *Cfb, Cd55*), cytokines (*Cxcl9, Il18rap*), and Jak/Stat pathway activation (*Stat1*, *Nfkb1*) (**Table S1**). Further, Module 6 expression on a per-tumor basis was strongly correlated with the level of T cell infiltration in the tumor, suggesting that it is a spatial signature of immune cell-mediated killing (**Figures S4B and C**). Taken together, these spatial gene expression data suggest that a low-level inflammatory immune reaction is a general feature of KRAS-driven tumor growth that is exacerbated by expression of *H3^K36M^* but is wholly dependent on SETD2 in both contexts.

### NK cells are sufficient for H3^K36M^-promoted tumor cell killing

In an orthogonal approach, to study the effects of H3^K36M^ expression on the immune response, we modified a *Kras*^G12D/+^; *Trp53^-/-^* mouse lung adenocarcinoma cell line (LG1233) [31] to express either *H3^WT^* or *H3^K36M^*, as well as an sgRNA targeting either *Setd2* or a non-targeting control to create four experimental groups: *KP-H3^WT^-Ctrl*, *KP-H3^WT^-Setd2^KO^*, *KP-H3^K36M^-Ctrl*, and *KP-H3^K36M^-Setd2^KO^* (**Figure 3A**). Importantly, all four cell lines had similar *in vitro* growth rates (**Figure S5A**). However, after orthotopic transplant of these cell lines into wildtype syngeneic mice, *KP-H3^K36M^-Ctrl* cells generated very small and diffuse tumors that were highly infiltrated with immune cells (**Figures S5B-E**). In contrast, similar to the autochthonous model, *KP-H3^K36M^-Setd2^KO^* cells robustly engrafted and expanded rapidly in the lungs of recipient mice. Thus, here again, the anti-tumor effect of *H3^K36M^* expression is dependent on *Setd2*.

**Figure 3:**
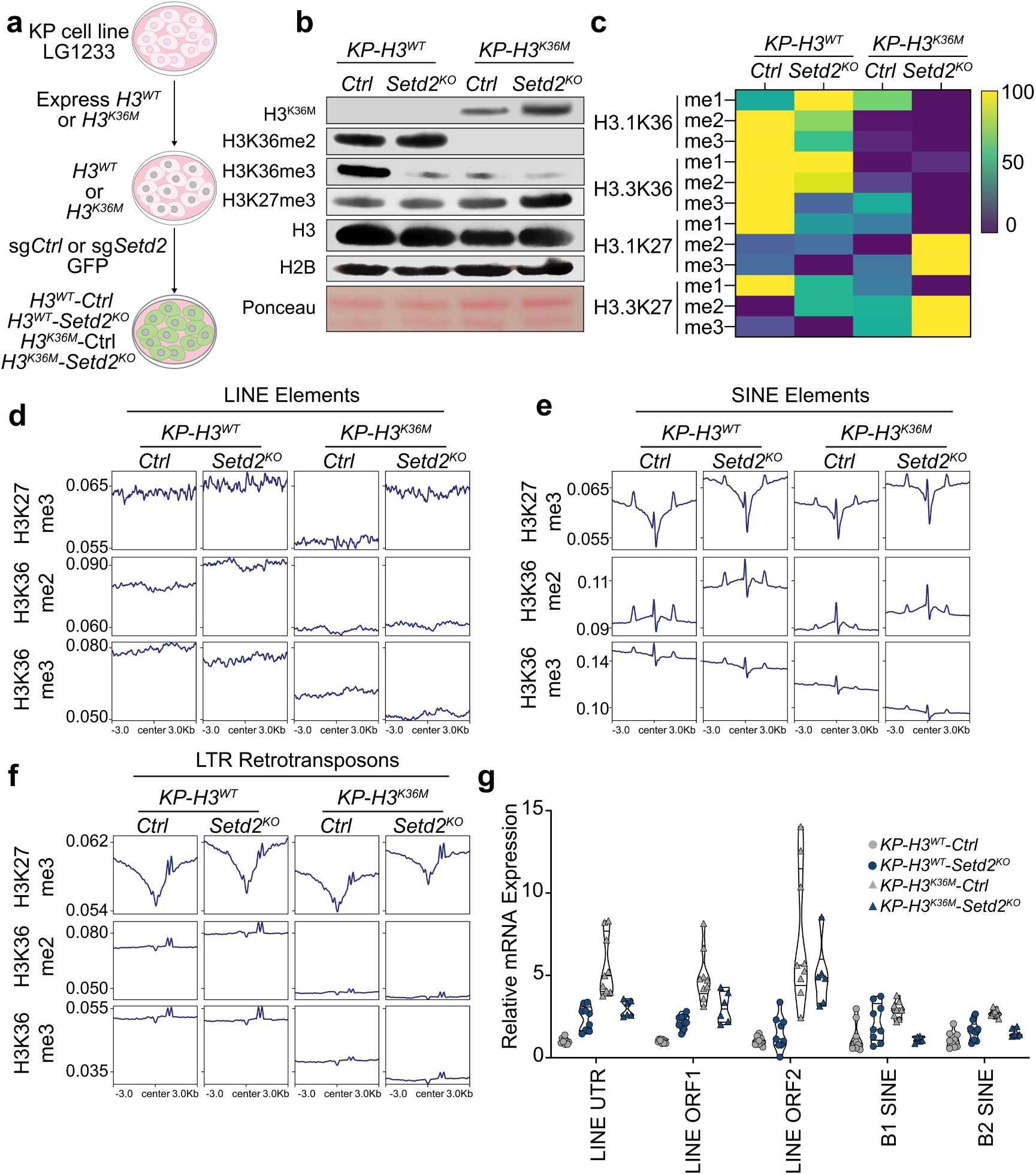
H3^K36M^ causes derepression of endogenous retroviral elements. a. Schematic for generation of *KP-H3^WT^-Ctrl*, *KP-H3^WT^-Setd2^KO^*, *KP-H3^K36M^-Ctrl*, and *KP-H3^K36M^-Setd2^KO^*cell lines. b. Immunoblot analysis of H3^K36M^, H3K36me2, H3K36me3, H3K27me3, H3, and H2B in histones prepared from *KP-H3^WT^-Ctrl*, *KP-H3^WT^-Setd2^KO^*, *KP-H3^K36M^-Ctrl*, or *KP-H3^K36M^-Setd2^KO^* cells. H3 and H2B are loading controls. c. Relative abundance of H3K36 and H3K27 histone modifications in *KP-H3^WT^-Ctrl*, *KP-H3^WT^-Setd2^KO^*, *KP-H3^K36M^-Ctrl*, and *KP-H3^K36M^-Setd2^KO^*cells by LC-MS/MS. Data are visualized in a row-normalized heat map. d. Metagene plot of ChIP-seq data for H3K27me3, H3K36me2, and H3K36me3 at LINE elements. Plots are centered on these elements across a ± 3 kb window. e. Metagene plot of ChIP-seq data for H3K27me3, H3K36me2, and H3K36me3 at SINE elements. Plots are centered on these elements across a ± 3 kb window. f. Metagene plot of ChIP-seq data for H3K27me3, H3K36me2, and H3K36me3 at LTR retrotransposons. Plots are centered on these elements across a ± 3 kb window. g. Quantitative real-time PCR for LINE and SINE element expression. Element expression is normalized to m5srRNA expression and is relative to expression of these elements in *KP-H3^WT^-Ctrl* cell lines.

To assess the necessity of an intact immune system on the subsequent clearance of *H3^K36M^*-expressing tumor cells, we orthotopically injected *KP-H3^WT^-Ctrl*, *KP-H3^WT^-Setd2^KO^*, *KP-H3^K36M^-Ctrl*, and *KP-H3^K36M^-Setd2^KO^* cell lines into triple-immunodeficient NOD-*Prkdc^em26Cd52^*;*Il2rg^em26Cd22^*(NCG) mice, which lack T, B, and NK cells. In this severely immunocompromised setting, all cell lines, including *KP-H3^K36M^* cells, form lung tumors at a similar efficiency (**Figures S6A and S6B**). However, when injected into athymic nude mice, which lack mature T cells, *KP-H3^K36M-^ Ctrl* cells were largely unable to form tumors, and those that did resulted in small tumors that were infiltrated with innate immune cells that expressed CD49b, which effectively marks NK cells in nude mice (**Figures S6C-S6E**). Concordantly, in an autochthonous setting, the NK-proficient, T and B cell-deficient *Rag1^KO^*background did not restore tumor growth of *KH3^K36M^*-expressing tumors (**Figures S6F-H**). Therefore, these data suggest that NK cells are sufficient to mediate the clearance of *H3^K36M^*-expressing tumor cells even in the absence of T cells.

### H3^K36M^ derepresses endogenous retroviral elements

To gain insight into cancer cell autonomous programs that are controlled by H3^K36M^ or SETD2, we performed RNA– and ATAC-sequencing on *KP-H3^WT^-Ctrl*, *KP-H3^WT^-Setd2^KO^*, *KP-H3^K36M^-Ctrl*, and *KP-H3^K36M^-Setd2^KO^* cells. Interestingly, while there were very few gene expression changes caused by *Setd2* inactivation in *H3^WT^*-expressing cell lines (*KP-H3^WT^-Ctrl* vs *KP-H3^WT^-Setd2^KO^*), *H3^K36M^*^-^expression induced gene expression changes that are typical of programs involved with an inflammatory immune response that include STAT signaling pathways and complement pathway activation (**Figures S7A and S7B**). Importantly, enrichment of these and other programs induced by *H3^K36M^* expression is dependent on *Setd2,* as they are significantly de-enriched in *KP-H3^K36M^-Setd2^KO^*cells compared to *KP-H3^K36M^-Ctrl cells* (**Figure S7B**). In contrast to the RNA-sequencing results, the ATAC-sequencing revealed very few changes in chromatin accessibility across groups, with cell lines clustering based on their H3^WT^ or H3^K36M^ status and *Setd2* inactivation having little to no effect (**Figure S7C**).

To elucidate the mechanism by which *H3^K36M^* expression promotes inflammation, we first measured global lysine methylation at histone tails via immunoblot and mass spectrometry on isolated histones from *KP-H3^WT^-Ctrl*, *KP-H3^WT^-Setd2^KO^*, *KP-H3^K36M^-Ctrl*, and *KP-H3^K36M^-Setd2^KO^*cells. As expected, relative to *KP-H3^WT^-Ctrl*, *KP-H3^WT^-Setd2^KO^*cells had reduced H3K36me3 while *H3^K36M^*-expressing cells had significantly reduced H3K36me3 and H3K36me2 (**Figures 3B, 3C, and S5F**). In contrast, global H3K27me3, which is generally associated with transcriptional repression, was significantly enriched in *KP-H3^K36M^-Setd2^KO^* cells. To examine positional changes in histone methylation across the genome, we performed ChIP-sequencing on *KP-H3^WT^-Ctrl*, *KP-H3^WT^-Setd2^KO^*, *KP-H3^K36M^-Ctrl*, and *KP-H3^K36M^-Setd2^KO^* cells. At gene bodies, *H3^K36M^* expression caused the depletion of H3K36me3, while *Setd2* inactivation had a more subtle effect (**Figure S7D**). *H3^K36M^* expression caused H3K36me2 enrichment primarily at transcription start sites (TSS) in *KP-H3^K36M^-Ctrl* cells and more broadly across gene bodies in *KP-H3^K36M^-Setd2^KO^* cells. In contrast, H3K27me3 deposition is complex but generally reduced across gene bodies in *H3^K36M^*-expressing cells and marginally affected by *Setd2* inactivation.

However, these complex patterns at gene bodies contrast with more salient patterns at endogenous retroviral elements (**Figures 3D-F**). Both H3K36me2 and H3K36me3 were significantly repressed by *H3^K36M^* expression at LINEs, SINEs, and LTR-retroelements. *Setd2* inactivation had subtle effects on H3K36me2 and H3K36me3 at LINEs, SINEs, and LTR-retroelements but minimally promoted H3K36me2 levels in *KP-H3^WT^-Setd2^KO^* and further repressed H3K36me3 in *KP-H3^K36M^-Setd2^KO^* cells. However, strikingly, H3K27me3 was significantly de-enriched across LINEs, SINEs, and LTR retroelements in *KP-H3^K36M^-Ctrl* cells. Moreover, this de-enrichment was fully dependent on *Setd2* and was restored in *KP-H3^K36M^-Setd2^KO^* cells. To determine if decreased H3K27me3 is associated with increased expression of elements, we performed RT-qPCR for LINE and SINE transcripts (**Figure 3G**). Element expression was highest in *KP-H3^K36M^-Ctrl* cells but significantly reduced in *KP-H3^K36M^-Setd2^KO^* cells. Taken together, these results suggest that expression of *H3^K36M^* promotes the derepression of retroviral elements by remodeling H3 methylation in a S*etd2-*dependent manner.

### H3^K36M^ promotes *Setd2*-dependent dsRNA synthesis

Changes in chromatin structure or function surrounding retroelements and other repeat sequences have been mechanistically linked to immune surveillance through the aberrant production or processing of RNA transcripts that ultimately form long double-stranded structures, or dsRNA [32]. Therefore, we hypothesized that the increased expression of retroviral transcripts caused by *H3^K36M^* expression could lead to the formation of dsRNA. Indeed, autochthonous *KH3^K36M^*lung tumors had significantly elevated levels of dsRNA compared to *K-Ctrl* tumors (**Figures 4A and 4B**). Additionally, elevated levels of dsRNA associated with *H3^K36M^*expression were dependent on *Setd2,* as the detection of dsRNA was lost in *KH3^K36M^-Setd2^KO^* tumors. Moreover, *KP-H3^K36M^-Ctrl* cells also had high levels of dsRNA, which was detected as multifocal puncta in the cytoplasm (**Figure 4C**). Here again, detection of dsRNA was minimal in *KP-H3^K36M^-Setd2^KO^* cells. Detection of dsRNA was confirmed using three independent dsRNA-specific antibodies, and specificity was confirmed by pretreatment with RNase III or transfection with synthetic dsRNA (**Figure 4C and S8**). In addition, *KP-H3^K36M^-Ctrl* cells synthesized high levels of IFN-β, a key interferon induced by dsRNA sensing, whereas *KP-H3^K36M^-Setd2^KO^*produced very low levels of this cytokine (**Figure 4D**). These data indicate that *H3^K36M^*-expression promotes cytoplasmic dsRNA accumulation in a *Setd2-*dependent manner.

**Figure 4:**
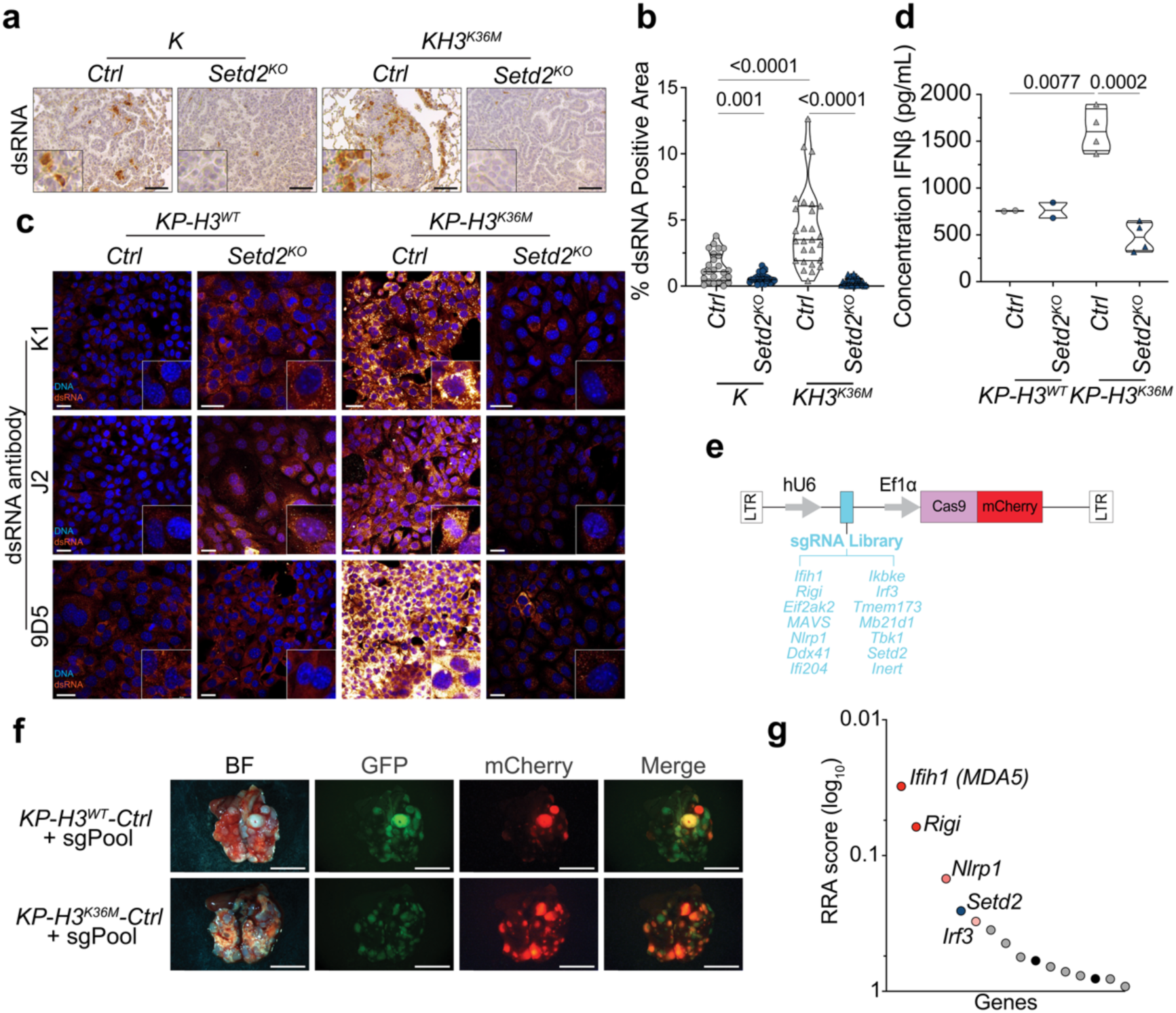
H3K36M promotes *Setd2*-dependent dsRNA synthesis. a. IHC for dsRNA in *K-Ctrl*, *K-Setd2^KO^*, *KH3^K36M^-Ctrl*, and *KH3^K36M^-Setd2^KO^* mice. Scale bars are 119 μm; insets are magnified x3. b. Quantification of dsRNA IHC in *K-Ctrl*, *K-Setd2^KO^*, *KH3^K36M^-Ctrl*, and *KH3^K36M^-Setd2^KO^* tumors. Significance was determined by unpaired Student’s t-test. Error bars represent mean±SEM. c. Immunocytochemistry for dsRNA in *KP-H3^WT^-Ctrl*, *KP-H3^WT^-Setd2^KO^*, *KP-H3^K36M^-Ctrl*, and *KP-H3^K36M^-Setd2^KO^* cells. Sparallel using 3 dsRNA antibodies: K1, J2, and 9D5. Scale bars are 25 μm. d. IFN-β ELISA analysis of cell culture supernatant collected from *KP-H3^WT^-Ctrl*, *KP-H3^WT^-Setd2^KO^*, *KP-H3^K36M^-Ctrl*, *KP-H3^K36M^-Setd2^KO^,* and cells. Significance was determined by unpaired Student’s t-test. Error bars represent mean±SEM. e. Schematic of plasmid used for CRISPR enrichment screen targeting dsRNA sensing pathway genes. f. Brightfield and fluorescent micrographs of lungs from athymic nude mice injected with *KP-H3^WT^-Ctrl* or *KP-H3^K36M^-Ctrl* cells infected with lentiCRISPRv2mCherry + sgRNA library. GFP marks all tumor cells while mCherry marks infected cells. Scale bars are 4.4 mm. g. Top-ranked genes enriched in CRISPR screen determined by comparing *KP-H3^K36M^-Ctrl* cells with *KP-H3^WT^-Ctrl* cells. Enriched genes are marked in red. Positive control *Setd2* is marked in blue. Non-targeting controls are marked in black.

To determine whether dsRNA expression and recognition are causal for the tumor-cell-killing immune responses observed, we employed a CRISPR-based screening approach targeting the key detectors of distinct aberrant nucleic acid structures, including dsRNA, or downstream pathway effectors that stimulate immune surveillance. We reasoned that inactivation of the critical detection factor(s) or signaling effector(s) would allow oncohistone-expressing cells (*KP-H3^K36M^-Ctrl*) to engraft and expand into tumors when transplanted into nude mice, where they would otherwise be rejected. Thus, we generated a CRISPR-based lentiviral transduction library consisting of three sgRNAs targeting each gene that encodes: STING, IFI16, DDX41, cGAS, MDA5, RIG-I, MAVS, IKKe, IRF3, and TBK1. In addition, five control sgRNAs that target safe harbor loci were included, as well as three sgRNAs targeting *Setd2* as a positively enriching control. These sgRNA-expressing constructs also express mCherry to track transduced cells (**Figure 4E**). Pooling all transduction vectors together, we transduced *KP-H3^WT^* or *KP-H3^K36M^* cells at a transduction multiplicity of ∼0.4. Each transduced cell population was orthotopically transplanted into the lungs of athymic nude mice and allowed to engraft and expand for 3 weeks. As expected, mice that received *KP-H3^WT^-Ctrl* cells had a mixture of both mCherry-positive and mCherry-negative tumors. In contrast, mice injected with *KP-H3^K36M^-Ctrl* cells had almost exclusively mCherry-positive tumors, indicating that transduced cells were preferentially selected during engraftment and expansion due to their ability to evade the anti-tumor immune response (**Figure 4F**). To determine the identity of the relevant genes whose inactivation drove selection, we extracted DNA from the tumor-bearing lungs, PCR amplified the sgRNA containing sequences and subjected the resulting amplicons to high-throughput sequencing. Compared to the relative sgRNA-specific read counts in lungs transplanted with *KP-H3^WT^-Ctrl* cells, the most enriched sgRNAs in the *KP-H3^K36M^-Ctrl* tumors targeted *Ifih1* (encoding MDA5) and *Rigi* (encoding RIG-I). MDA5 and RIG-I together make up the core components of the major dsRNA sensing pathway traditionally associated with antiviral mechanisms (**Figure 4G**). As expected, *Setd2*-targeting sgRNAs were also highly enriched, and sgRNAs targeting *Nlrp1* and *Irf3* were also enriched to a lesser extent. These findings indicate that the ability of the cell to detect dsRNA species is critical for tumor-cell-killing immune stimulation induced by *H3^K36M^* and again demonstrate the critical requirement for *Setd2* in the process.

### Inactivation of H3K36 di-methyltransferases phenocopies H3^K36M^ expression

The pleiotropic ability of oncohistones to alter deposition of both H3K36me2 and H3K36me3 across the genome suggests that other H3K36 methyltransferases beyond SETD2 may play a role in oncohistone-mediated dsRNA production and immune surveillance. Additionally, that oncohistone expression profoundly reduced H3K36me2 at retroelements suggests that dominant inhibition of di-methyltransferases, namely NSD1 or NSD2, may play a causal role in the derepression of retroelements and subsequent immune surveillance (**Figures 3D-F**). To determine whether *Nsd1* or *Nsd2* inactivation could replicate the effects of H3^K36M^ expression in autochthonous KRAS^G12D^-driven lung cancer, we used the same CRISPR-based somatic genome editing approach as outlined above (**Figure 5A**). Compared to *K-Ctrl* tumors, *K-Nsd1^KO^*and *K-Nsd2^KO^* tumors were significantly fewer in number and smaller in size (**Figures 5B, 5C, and S9A**). Indeed, *K-Nsd1^KO^* tumors were detected in only one mouse across all replicates, indicating that loss of NSD1 is largely incompatible with KRAS-driven lung tumor growth or that NSD1 is generally required for cell viability. In contrast, though less frequent than *K-Ctrl* tumors, *K-Nsd2^KO^*tumors were present in most replicate mice. *K-Nsd2^KO^* tumors had significantly diminished H3K36me2, highly elevated dsRNA, and were inundated with immune cells (**Figures S9B, 5D, and 5E**). Further, *KP-Nsd2^KO^* cell lines produced high levels of IFN-β (**Figures 9C and 9D**). Thus, essentially mirroring the effects of *H3^K36M^* expression, these data suggest that the dsRNA-triggered, immune-mediated, and anti-tumor effects are likely due to oncohistone-mediated antagonism of NSD2 and nominate NSD2 as a target for the treatment of KRAS-driven lung adenocarcinoma.

**Figure 5:**
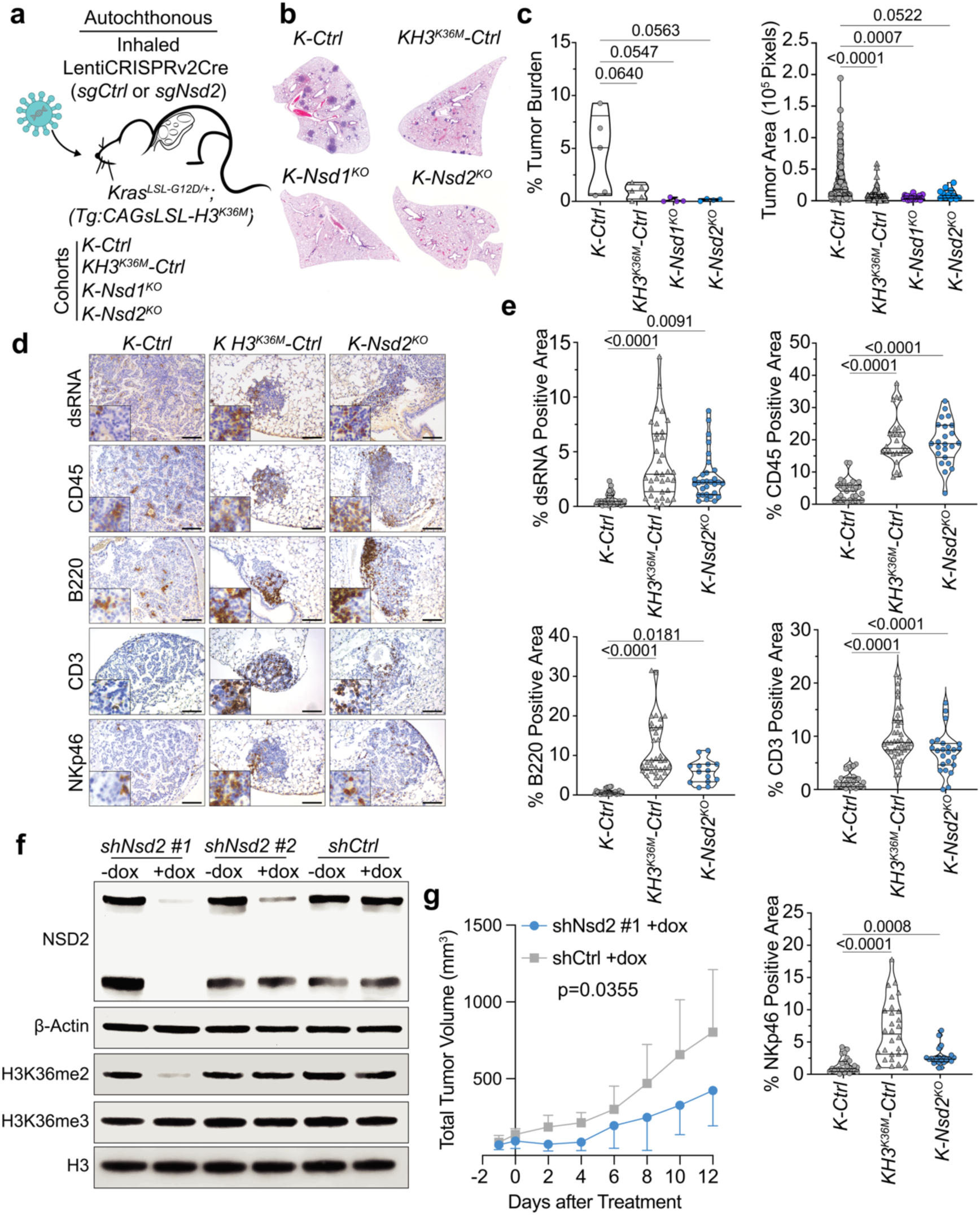
Inactivation of H3K36 di-methyltransferase *Nsd2* phenocopies H3^K36M^ expression. a. Experimental scheme of autochthonous tumor initiation. b. Representative images of tumor bearing lobes from *K-Ctrl*, *KH3^K36M^-Ctrl*, and *K-Nsd2^KO^* mice. c. Quantification of tumor burden and area in *K-Ctrl*, *KH3^K36M^-Ctrl*, and *K-Nsd2^KO^* mice. Significance was determined by unpaired Student’s t-test. Error bars represent mean±SEM. d. IHC for dsRNA and immune cell markers CD45, B220, CD3, and NKp46 in *K-Ctrl*, *KH3^K36M^-Ctrl*, and *K-Nsd2^KO^* tumors. Scale bars are 119 μm; insets are magnified x3. e. Quantification of dsRNA, CD45, B220, CD3, and NKp46 IHC staining in *K-Ctrl*, *KH3^K36M^-Ctrl*, and *K-Nsd2^KO^* mice. Significance was determined by unpaired Student’s t-test. Error bars represent mean±SEM. f. Immunoblot analysis of NSD2 and B-Actin in whole cell lysates and analysis of H3K36me2, H3K36me3, and H3 in histones prepared from *KP-*shNsd2 *#1*, *KP-*shNsd2 *#2*, and *KP-*shCtrl cells with and without dox treatment after 48 hours. g. Tumor growth curves for *KP-*shNsd2 #1 and *KP-*shCtrl cells injected into male C57BL/6 mice. Significance determined by two-way ANOVA test. Error bars represent mean±SEM.

To model the potential therapeutic benefits of NSD2 inhibition in established KRAS*-*driven lung adenocarcinoma tumors, we engineered *KP* cells with doxycycline-inducible shRNAs targeting *Nsd2* (**Figures S9E and S9F**). GFP expression is coupled with shRNA expression to mark *Nsd2* silencing. We tested two distinct shRNAs targeting *Nsd2* and found one that strongly reduced NSD2 protein expression and downstream H3K36me2 upon doxycycline treatment, but did not affect H3K36me3 levels (**Figures 5F and S9G**). We subcutaneously injected this cell line or a doxycycline-inducible shCtrl cell line into the flanks of immunocompetent mice. After the tumors reached an average size of 100mm^3^, we administered mouse chow containing 625mg/kg of doxycycline to induce shRNA expression. Compared to tumors expressing shCtrl that grew steadily, expression of *Nsd2* knockdown resulted in acute tumor regressions.

However, each tumor ultimately escaped *Nsd2* knockdown and expanded, albeit at a slower rate compared to shCtrl expressing tumors (**Figure 5G**). To investigate the effect of *NSD2* expression in human lung adenocarcinoma, we performed gene set enrichment analysis (GSEA) on patient lung adenocarcinoma gene expression data. Consistently, gene sets related to T and NK cell function were downregulated in patients with high *NSD2* expression (**Figure S9H**). Further, we found that low *NSD2* expression is associated with significantly increased overall survival in lung adenocarcinoma patients (**Figure S9I**). Taken together, these results suggest that H3^K36M^ promotes tumor-cell-killing via inhibition of NSD2 and support the pursuit of NSD2 inhibitors as a therapeutic modality in KRAS-driven lung adenocarcinoma.

## DISCUSSION

Despite the bona fide oncogenic potential of H3^K36M^ oncohistone in many tumor types, our data clearly demonstrate its paradoxical effect as a tumor suppressor in KRAS-driven lung adenocarcinoma. Moreover, that H3^K36M^ mediates tumor suppression while limiting the actions of SETD2, the only identified mammalian H3K36 tri-methyltransferase and a potent tumor suppressor in KRAS-driven lung adenocarcinoma, is doubly surprising. Our data demonstrate that the complex mechanism of action for H3^K36M^ in dominantly inhibiting multiple H3K36 methyltransferases underpins this dichotomy. Indeed, we show that inactivation of the major H3K36 dimethyltransferase *Nsd2* phenocopies *H3^K36M^* expression to promote derepression of endogenous retroviral element transcription, formation of dsRNA, and induction of an antiviral-like immune response that severely limits tumor growth *in vivo* (**Figure S10**). Perhaps most surprising is the contrasting effect of *Setd2* inactivation in the context of oncohistone expression. Though demonstrated to be a potent tumor suppressor in the context of oncogenic KRAS [7, 8, 10, 33], that SETD2 inactivation completely reverses the anti-tumor effects of oncohistone expression was completely unexpected. This is made more surprising by the expectation that oncohistone expression would functionally inactivate SETD2 due to its demonstrated dominant mechanism of action [2, 3, 34]. Though our data show that oncohistone expression in KRAS-driven tumors significantly limits H3K36me3, clearly any remaining activity of SETD2 is sufficient to allow or even facilitate the potent antiviral-like, tumor-eradicating immune response observed. It is tempting to speculate that SETD2 has additional roles beyond tri-methylation of H3K36 that are not suppressed by H3^K36M^ and may explain its immune-regulating effects. Indeed, SETD2 is reported to directly methylate and promote phosphorylation/activation of STAT1, which is a major transcriptional mediator of antiviral immune responses [21]. Additionally, SETD2 was demonstrated to regulate PRC2-dependent H3K27me3 by directly methylating and thereby promoting EZH2 degradation [22]. Thus, in the context of oncohistone expression that elicits a chromatin-derepression-based mechanism which ultimately provokes an antiviral-like immune-mediated anti-tumor response, SETD2 may be the master regulator through these non-histone-directed mechanisms. However, it remains deeply curious that the cellular decision to induce non-cell autonomous death or to proliferate uncontrollably happens via the opposing actions of distinct H3K36 methyltransferases. Finally, our data implicates the inactivation of *Nsd2* as a stimulator of immune surveillance. Translation of these findings into clinically actionable modalities could be important for cancer treatment.

## METHODS

### Ethics statement

The research in this study was conducted ethically and complies with all relevant guidelines and regulations. Animal studies were performed under strict compliance with the Institutional Animal Care and Use Committee (IACUC) at the University of Pennsylvania (#804774).

### Animal studies and treatment

For the autochthonous lung adenocarcinoma mouse model, *Kras^LSL-G12D^* mice (Jax stock number 008179), *Rosa26^LSL-YFP/LSL-YFP^*, and *LSL-H3^K36M^* transgenic mice have been previously described [35–37]. Mice are C57BL/6J. Mice were transduced with 6 x 10^4^ plaque-forming units (PFUs) per mouse of lentiCRISPRv2Cre by endotracheal intubation [38]. LentiCRISPRv2Cre expressing sgRNAs targeting *Setd2*, *Nsd2,* and β*-Galactosidase (βgal,* labeled as *sgCtrl*) were used. The sgRNA sequences are listed in **Table S2**. *sgSetd2* and *sgCtrl* have been described previously [8, 39, 40]. *sgNsd2* was designed as described below in “CRISPR design and production”. Lentivirus production and titration were performed as described previously using HEK293B cells [8].

For the orthotropic lung adenocarcinoma mouse model, *Rosa26-LSL-Cas9* mice (called C57BL/6J mice, Jax stock number 026175), OD-Prkdc^em26Cd52^; Il2rg^em26Cd22^ mice (NCG, Taconic strain 572), and CrTac: NCr-*Foxn1nu* mice (athymic nude, Taconic strain NCRNU) have been previously described. 50,000 cells were injected into the tail veins of these mice at ∼4-5 weeks of age. For C57BL/6J, NCG, and athymic nude mice, tumors formed for 5, 2, or 3 weeks, respectively. Only male NCG and athymic nude mice were used. For all other mouse experiments, roughly equal proportions of male and female mice were used. The animal protocol was approved by the University Laboratory Animal Resources (ULAR) at the University of Pennsylvania and the IACUC. Fluorescent images of GFP-or mCherry-positive tumors in mouse lungs were captured using a Leica M80 compact stereomicroscope.

In the subcutaneous tumor experiment, 100,000 *KP-*shNsd2 or *KP-*shCtrl cells were subcutaneously injected into the flanks of *Rosa26-LSL-Cas9* mice (called C57BL/6J mice, Jax stock number 026175). Tumor volumes were measured prior to the start of doxycycline treatment, and the mice were put on a doxycycline diet (Envigo Teklad, TD.01306) after the average tumor size reached 100mm^3^. Tumor volumes were then measured every two days following treatment. Tumor volumes were calculated using the formula: 0.52 x a x b^2^ where a is the larger diameter and b is the smaller diameter of the tumor mass. All mice were euthanized once the average tumor volume was 2000 mm^3^.

### Immunohistochemistry and Immunofluorescence

Lung and tumor tissue were dissected into 10% neutral-buffered formalin overnight at room temperature before paraffin embedding. Paraffin-embedded and H&E histological sections were produced by the Penn Molecular Pathology and Imaging Core. Immunostaining for H3^K36M^ (1:2500, Cayman Chemical Company 31547), Ki67 (1:800, Novus Biologicals NB110-89717), pH3 (1:500, Cell Signaling Technology cs9701), Nkx2.1 (abcam ab76013), and CD3 (1:700, abcam ab5690) was performed using the Leica BOND RXm Automatic Slide Stainer. Immunostaining for SPC (1:2000, Millipore AB3786), CD45 (1:100, BioLegend 103102), B220 (1:50, BD Biosciences 551459), NKp46 (1:100, BioLegend 137602), F4/80 (1:500, Bio-Rad MCA497GA), Ly6G (1:500 Biolegend), and CD49b (1:500, Thermo Fisher 14-5971-85) was performed after deparrafinization and citrate-based antigen retrieval. Immunostaining for H3K36me3 (1:2000, abcam ab9050) and H3K36me2 (1:1000, abcam ab9049) was performed after proteinase K antigen retrieval. Formalin-fixed, paraffin-embedded sections were incubated with avidin and biotin blocking steps for 20 minutes each (Vector Laboratories, SP-2001), a 30-minute protein block (Dako, X090930-2), primary antibody overnight at 4 °C, secondary antibody for 1 h at room temperature, and with ABC reagent for 30 min at room temp (Vector Laboratories PK-4001 for rabbit antibodies and Vector Laboratories PK-6104 for rat antibodies). For J2 dsRNA (1:500, Cell Signaling Technology 76651) immunostaining, citrate-based antigen retrieval was performed, and then a Mouse on Mouse Immunodetection kit was used (Vector Laboratories, BMK-2202) according to manufacturer’s instructions.

For TUNEL staining, tissues were deparaffinized and then permeabilized with 0.1% sodium citrate and 0.1% Triton-X in PBS for 8 min. TMR Red-conjugated TUNEL labeling mix (Millipore Sigma, 12156792910) was added to tissue sections and incubated for 1 h at 37 °C in a humidity chamber. Nuclei were stained by mounting the slides with DAPI-containing Fluoro-Gel (Electron Microscopy Sciences, 17985-50). All photomicrographs were captured on a Leica DMI6000B inverted light and fluorescence microscope.

### Histological quantification

ImageJ software was used to quantify immunostained samples. Quantification was carried out as previously described [41] for immune markers, Ki67, and J2 dsRNA. For TUNEL quantification, positive cells were counted and divided by the total area of the tumor.

### Analysis of Xenium Spatial Transcriptomics data

Cell typing was performed using Scanpy. Briefly, low-quality cells (<55 unique genes) were filtered out, and counts were normalized to cell area. The k-nearest neighbors graph was computed with k=10 and n=35 PCs, after which cells were embedded with UMAP and coarsely clustered at resolution 0.5 with Leiden clustering. Clusters were annotated based on differentially expressed genes and, in some cases, spatial distributions. For instance, AT2 cells and AT2-like malignant cells initially clustered together due to similar gene expression profiles but could be distinguished in sub-clustering by spatial location (ie, in tumor nodules or lung parenchyma). Cellular neighborhood analysis was performed using Scimap. First, the nearest 30 spatial neighbors were determined for each cell, and cells were k-means clustered, using k=30 to overcluster cells into neighborhoods. Then, neighborhoods were manually merged based on similarity in cell type abundance and distribution in space. Individual tumors were identified by computing the density of malignant cells in 50 x 50 micron bins, and bins were marked as ‘tumor’ if they contained >20% malignant cells. Tumors were further pruned if they had fewer than 10 malignant cells, resulting in a total of 159 tumors across the four genotypes. T cell activation score was computed using Scanpy’s score_genes function on the gene list GO:0050870 (Positive regulation of T cell activation), considering only genes measured in the Xenium experiment.

### Cell Lines

LG1233 cells were generated from lung tumors of C57BL/J6 *Kras*^G12D/+^; *Trp53^flox/flox^* mice and kindly provided by Dr. Tyler Jacks (Massachusetts Institute of Technology, Cambridge, USA). All cell lines were grown in high-glucose DMEM supplemented with 10% fetal bovine serum, GlutaMAX, and gentamicin at 37 °C and 5% CO_2_. *H3.3^WT^* or *H3.3^K36M^* genes were cloned into MSCV puro (Addgene 68469). Retrovirus was produced, and LG1233 cells were infected using polybrene at 4μg/ml. Infected cells were selected using 2.5 μg/ml puromycin. Next, lentivirus was produced using lentiCRISPRv2GFP (Addgene 82416) with sgRNAs targeting either *Setd2* or *βgal* (sgRNA sequences above and in **Table S2**). *KP-H3^WT^* or *KP-H3^K36M^*cells were infected with these lentiviruses and then sorted for GFP positivity by FACS. Next, 1000 resultant cells were plated on a 15cm dish and expanded for 1-2 weeks until colonies formed. Colonies were individually trypsinized and grown separately as clones. Clones were then screened for *H3^K36M^* expression and *Setd2^KO^*by immunoblot analysis.

Cell number was counted using a Beckman Coulter Z2 Cell and Particle Counter. Proliferation of cell lines was determined by plating 25,000 cells, counting cells, and replating 25,000 cells every 4 days in triplicate.

Nsd2 shRNA sequences were cloned into the doxycycline-inducible lentiviral plasmid pCF806-recI (Addgene 186711). shRNA sequences were designed using VectorBuilder’s shRNA Target Design tool (https://en.vectorbuilder.com/tool/shrna-target-design.html). The plasmid pCF806-shRen.713 (Addgene 186712) was used as a nontargeting control. Briefly, shRNA 97mer oligos (sequences in **Table S3**) were PCR amplified using miRE_F: TGAACTCGAGAAGGTATATTGCTGTTGACAGTGAGCG and miRE_R: GCTCGAATTCTAGCCCCTTGAAGTCCGAGG primers. PCR products were then ligated into XhoI/EcoRI-digested pCF806-recI (NEB, M0202S). Lentivirus production was performed, and LG1233 cells were transduced. Knockdown was confirmed by immunoblot analysis for NSD2 and H3K36me2.

### Immunoblot analysis

For whole cell lysates, cells were lysed in RIPA buffer. Acid-extracted histones were prepared for histone methylation Western blots. Samples were resolved on NuPage 4–12% (for whole cell lysates) or 12% (for histones) Bis-Tris protein gels (Thermo Fisher) and transferred to polyvinylidene fluoride (PVDF) membranes. Blocking, primary, and secondary antibody incubations were performed in Tris-buffered saline (TBS) with 0.1% Tween-20. Histones were blotted for H3^K36M^ (1:1000, Cayman Chemical Company 31547), H3K36me3 (1:1000, Abcam ab9050), H3K36me2 (1:1000 Abcam ab9049), H3K27me3 (1:1000, Cell Signaling Technology 9733), H3 (1:20000, Abcam ab1791), and H2B (1:20000, Abcam ab1790). Whole cell lysates were blotted for NSD2 (1:1000, Abcam, ab75359) and β-actin (1:10000, Sigma Aldrich, A2066). Protein concentration was determined using a Bradford Protein Assay Kit (Thermo Scientific A55866).

### Histone extraction

Histone proteins were extracted from the pellet as described by [42] to ensure good-quality identification and quantification of single histone marks. Briefly, histones were acid-extracted with 0.4 N sulfuric acid and incubated with constant rotation for 30 min, followed by precipitation with 33% trichloroacetic acid (TCA) for 1 h at 4C. Then, the supernatant was removed and the tubes were rinsed with acetone containing 0.1% 12N HCl, centrifuged, and rinsed again using 100% acetone. After the final centrifugation, the supernatant was discarded, the pellet was dried at room temperature for 30 min, and the histones were resuspended in molecular-grade H_2_O. The histones were then used for either immunoblotting or mass spectrometry.

### Histone digestion and desalting

The pellet was dissolved in 50 mM ammonium bicarbonate, pH 8.0, and histones were subjected to derivatization using 5 µL of propionic anhydride and 14 µL of ammonium hydroxide (all Sigma Aldrich) to balance the pH at 8.0. The mixture was incubated for 15 min and the procedure was repeated. Histones were then digested with 1 µg of sequencing grade trypsin (Promega) diluted in 50mM ammonium bicarbonate (1:20, enzyme:sample) overnight at room temperature. Derivatization reaction was repeated to derivatize peptide N-termini. The samples were dried in a vacuum centrifuge. Prior to mass spectrometry analysis, samples were desalted using a 96-well plate filter (Orochem) packed with 1 mg of Oasis HLB C-18 resin (Waters). Briefly, the samples were resuspended in 100 µl of 0.1% TFA and loaded onto the HLB resin, which was previously equilibrated using 100 µl of the same buffer. After washing with 100 µl of 0.1% TFA, the samples were eluted with a buffer containing 70 µl of 60% acetonitrile and 0.1% TFA and then dried in a vacuum centrifuge.

### LC-MS/MS Acquisition and Analysis

Samples were resuspended in 10 µl of 0.1% TFA and loaded onto a Dionex RSLC Ultimate 300 (Thermo Scientific), coupled online with an Orbitrap Fusion Lumos (Thermo Scientific). Chromatographic separation was performed with a two-column system, consisting of a C-18 trap cartridge (300 µm ID, 5 mm length) and a picofrit analytical column (75 µm ID, 25 cm length) packed in-house with reversed-phase Repro-Sil Pur C18-AQ 3 µm resin. Peptides were separated using a 30 min gradient from 1-30% buffer B (buffer A: 0.1% formic acid, buffer B: 80% acetonitrile + 0.1% formic acid) at a flow rate of 300 nl/min. The mass spectrometer was set to acquire spectra in a data-independent acquisition (DIA) mode. Briefly, the full MS scan was set to 300-1100 m/z in the orbitrap with a resolution of 120,000 (at 200 m/z) and an AGC target of 5×10e5. MS/MS was performed in the orbitrap with sequential isolation windows of 50 m/z with an AGC target of 2×10e5 and an HCD collision energy of 30.

Histone peptides raw files were imported into EpiProfile 2.0 software [43]. From the extracted ion chromatogram, the area under the curve was obtained and used to estimate the abundance of each peptide. In order to achieve the relative abundance of post-translational modifications (PTMs), the sum of all different modified forms of a histone peptide was considered as 100% and the area of the particular peptide was divided by the total area for that histone peptide in all of its modified forms. The relative ratio of two isobaric forms was estimated by averaging the ratio for each fragment ion with different mass between the two species. The resulting peptide lists generated by EpiProfile were exported to Microsoft Excel and further processed for a detailed analysis.

### RNA-sequencing Preparation and Analysis

RNA isolation, library preparation, and sequencing of 3 clonal cell lines per group (*KP-H3^WT^-Ctrl*, *KP-H3^WT^-Setd2^KO^*, *KP-3^K36M^-Ctrl*, and *KP-H3^K36M^-Setd2^KO^*) were performed by Genewiz/Azenta Life Sciences. Libraries were subjected to Illumina 150-bp paired-end sequencing.

Reads from fastq files were mapped to the mouse genome (GRCm39) using Salmon with ‘libType A’ and ‘validateMapping’ options [44]. The reads mapped to coding genes were then used as an input to DESeq to perform differential analysis [45]. For volcano plot data, differentially expressed genes were defined as genes with a false-discovery rate-adjusted P value less than 0.05 and log_2_-normalized fold change in expression greater than ±1. For the pathway enrichment analysis, the gene counts were used as input to the GSEA program [46]. Enrichment analysis was performed on the Mouse Hallmark and Biological Processes gene sets from MSigDb [46]. Heatmaps were generated using the normalized counts from DESeq using the pheatmap package [47]. In addition, ashr [48], ggplot [49], snakemake [50], tximport [51], and rtracklayer [52] packages were used for performing these analyses.

### ATAC-sequencing Preparation and Analysis

ATAC-seq libraries were generated and sequenced by Azenta Life Sciences from 3 clonal cell lines per group (*KP*-*H3^WT^-Ctrl*, *KP*-*H3^WT^-Setd2^KO^*, *KP*-*H3^K36M^-Ctrl*, and *KP*-*H3^K36M^-Setd2^KO^*). Libraries were subjected to Illumina 150-bp paired-end sequencing.

ATAC-seq FASTQ files were trimmed using Atria (default options) to remove adapter reads and were then aligned to mm39 (GRCm39) using bowtie2 (“bowtie align” using options –local –no-mixed –discordant, v2.5.1) [53]. Peak calling was performed on the aligned reads using Genrich with the –j and –d 150 options [54]. BigWig files were generated using bamCoverage from deeptools. Using deeptools bigWigsummary and plotCorrelation, the correlation heatmaps were plotted for these samples [55].

### Chromatin immunoprecipitation (ChIP)

15 million cells from 2 clonal cell lines per group (*KP-H3^WT^-Ctrl*, *KP-H3^WT^-Setd2^KO^*, *KP-H3^K36M^-Ctrl*, and *KP-H3^K36M^-Setd2^KO^*) were harvested and cross-linked at room temperature by resuspending cells in PBS containing 1% PFA for 5 min. Crosslinking was quenched by the addition of glycine to a final concentration of 125 mM. After washing cells twice with PBS, cells were lysed in 1 mL sonication buffer (0.25% Sarkosyl, 1 mM DTT, and protease inhibitor in RIPA buffer). Cell suspensions were sonicated using a Covaris S220 Focused Ultrasonicator until chromatin was sheared for 17 min per sample to a size range of around 200-500 bp. Lysates were centrifuged at a maximum speed for 5 min, supernatant was collected, and lysates were flash frozen in liquid nitrogen. The next day, Protein A Dynabeads (Invitrogen, 10001D) were incubated for 6 hours with H3K36me2 (5 µg, abcam, ab9049), H3K36me3 (5 µg, abcam, ab9050), H3K27me3 (5 µg, Cell Signaling Technologies, 9733T), or mouse IgG (1 µg, Diagenode, C15400001-15) antibody at 4°C. Sonicated chromatin was thawed and mixed with 2% sonicated S2 chromatin as a spike-in control. Chromatin from each sample was mixed and used as a single input control. Sonicated samples were added to the pre-bound Dynabeads and rotated overnight at 4°C. Chromatin-bound beads were washed three times with low-salt wash buffer (0.1% SDS, 1% Triton X-100, 1 mM EDTA pH 8.0, 50 mM Tris-HCl at pH 8.0, 150 mM NaCl), high-salt wash buffer (0.1% SDS, 1% Triton X-100, 1 mM EDTA pH 8.0, 50 mM Tris-HCl pH 8.0, 500 mM NaCl), and LiCl wash buffer (150 mM LiCl, 0.1% SDS, 0.5% deoxycholic acid sodium salt, 1% NP-40, 1 mM EDTA pH 8.0, 50 mM Tris pH 8.0). Beads were then washed with TE buffer containing 50 mM NaCl and spun down at 950g. The beads were resuspended in ChIP elution buffer (1% SDS, 200 mM NaCl, 10 mM EDTA, pH 8.0, 50 mM Tris pH 8.0) and eluted for 30 min at 65°C. The solutions were spun down for 1 min at 16000 g, and supernatants were transferred to new tubes. Input DNA was diluted to 200 µl using ChIP elution buffer. Input and experimental samples were reverse cross-linked overnight at 65°C. All samples were incubated with RNase A for 1 hour at 37°C, followed by Proteinase K treatment for 1 hour at 55°C. DNA was purified using a PCR purification Kit (Qiagen, 28104) and eluted into 50 µl of molecular biology grade water. Libraries were generated by Genewiz/Azenta Life Sciences (2 × 150 bp paired-end sequencing; Illumina HiSeq).

### ChIP sequencing analysis

For ChIP-seq samples, spike-in normalization was performed as in [56]. Drosophila chromatin was added to the histone ChIP samples before the IP was performed to a concentration of 2%. The reads from these samples were first trimmed using Atria (default options, v4.0.0) [57] and then aligned to mm39 (GRCm39) using bowtie2 (“bowtie align” using options –local –no-mixed – discordant, v2.5.1) [53]. Reads were then also aligned to the Drosophila genome dm6 (BDGP Release 6 + ISO1 MT), and the number of reads mapping to this genome was counted, and the percentage of Drosophila reads in each sample was calculated. Scaling factors for each sample were then calculated by scaling up human reads such that the percentage of Drosophila reads in each sample was the same. Using DeepTools [55], bigWig files for the samples were then generated using the scaleFactors option while providing respective scaling factors calculated in the previous step. RepeatMasker masked repeat Sequences were downloaded from the UCSC Genome browser database [58]. Using Deeptools, matrices were computed using the ‘computeMatrix’ command for the different repeat elements and plotted using the ‘plotHeatmap’ command.

### Quantitative reverse transcription-PCR

Total RNA was extracted from cells using Qiagen RNeasy Mini Kit (Qiagen, 74106). cDNA synthesis was performed using High-Capacity cDNA Reverse Transcription Kit (Applied Biosystems, 4368814). RT-PCR was performed using SYBR Green I Nucleic Acid Gel Stain (Invitrogen, S7563) in triplicate, following manufacturer’s instructions, and evaluated on an Applied Biosystems ViiA 7 RT-PCR machine. RT-PCR primers are listed in **Table S4**.

### Immunocytochemistry

25,000 *KP*-*H3^WT^-Ctrl*, *KP*-*H3^WT^-Setd2^KO^*, *KP*-*H3^K36M^-Ctrl*, and *KP*-*H3^K36M^-Setd2^KO^* cells were plated in chamber slides (EMD Millipore PEZGS0816) and grown overnight. Cells were then fixed with 4% PFA, permeabilized with 0.1% Triton X-100 detergent in 1X PBS, blocked with 1% BSA in 1X PBS, and stained overnight with anti-dsRNA antibodies K1 (1:800, Cell Signaling Technology 28764), J2 (1:500, Cell Signaling Technology 76651), or 9D5 (1:500, Ab00458-23.0) diluted in blocking buffer. Cells were then stained with either anti-rabbit IgG Alexa Fluor 594 (1:200, Thermo Fisher A32754) or anti-mouse biotinylated secondary antibody (1:200, Vector Laboratories BA-9200) for 1 hour at room temperature, and then with antistrepavidin Alexa Fluor 594 (1:200, Thermo Fisher S32356). Nuclei were stained with DAPI. Cells were then mounted with Fluoromount-G (Southern Biotech 0100-01), and imaging was performed on a Leica Stellaris 5 confocal microscope using a 63x oil objective.

For RNase III treatment, cells were treated with ShortCut RNase III (New England Biolabs M0245) after permeabilization for 2 hr at room temperature according to manufacturers instructions. For poly(I:C) treatment (Sigma Aldrich P1530), cells were plated as above and then transfected with 0.25 μg poly(I:C) using Lipofectamine 3000 Transfection Reagent (Thermo Fisher L3000001) following manufacturer’s instructions. For both of these treatments, the staining protocol was then followed as above.

### IFNB1 ELISA

Mouse interferon beta (IFNB1) levels were quantified using the Mouse IFNB1 ELISA Kit (Aviva Systems Biology, OKEH00314), according to manufacturer’s instructions. Briefly, pre-coated anti-IFNB1 plates were incubated with standards and experimental samples for 2 h at room temperature. 1x biotinylated IFNB1 detection antibody (1:100) was added to each well and incubated for 1 hour. Samples were then incubated with 1X Avidin-HRP conjugate (1:100) for 30 minutes. Lastly, TMB substrate was added to develop the colorimetric signal. The reaction was stopped with the provided stop solution, and absorbance was measured at 450 nm using a Tecan Infinite 200 Pro microplate reader. Sample concentrations were determined using the standard curve generated using serial dilutions of the IFNB1 standard.

### CRISPR design and production

All sgRNAs not delineated above were designed using the Sanjana Lab CRISPR Cas9 library design tool (http://guides.sanjanalab.org). For *in vivo* CRISPR, sgRNA duplexes were Golden Gate cloned into BsmBI sites of the LentiCRISPRv2-Cre vector. LentiCRISPRv2GFP sg*Setd2* and sg*Ctrl* were cloned in this manner as well. For the CRISPR enrichment screen library generation, the In-Fusion cloning system (Clontech: #638909) was used. The sgRNA pool was cloned by annealing two DNA oligos for each guide and ligating into a BsmB1-digested lentiCRISPRv2mCherry vector (Addgene 99154), as described [59]. The sgRNA sequences used are detailed in **Table S2**.

### Pooled CRISPR Enrichment Screen

The sgRNA library targeting dsRNA sensing pathway components was generated as described above. Lentivirus was produced and titered. *KP-H3^WT^-Ctrl* and *KP-H3^K36M^-Ctrl* cells were infected at an MOI of 0.4 to ensure a single copy of sgRNA transduction per cell. 50,000 of the resultant cells were injected into the tail veins of 10 athymic nude mice per group. After 3 weeks, the mice were euthanized, their lungs were isolated, and fluorescent images were taken of their lung tumors. Each lung tree was placed into 10 mL of digestion buffer containing 100 mM NaCl, 20 mM Tris, 10 mM EDTA, 0.5% SDS, and 100 μL of 20mg/mL Proteinase K. The tissue was cut into small pieces and incubated at 55°C overnight. The next day, genomic DNA was extracted from 5 mL of each digested sample by phenol-chloroform extraction.

To amplify the tumor sgRNA sequences, 20 μg of genomic DNA was PCR amplified. The genomic DNA from the original cell lines infected with the sgRNA library was also extracted and PCR amplified as a benchmark for sgRNA representation. Q5 Hot Start High-Fidelity 2x Master Mix (New England Biolabs, M0494X) was mixed with genomic DNA and a universal forward and reverse primer set (Forward: 5’ CGTGACGTAGAAAGTAATAATTTCTTGGGTAG 3’, Reverse: 5’ GAGGCCGAATTCAAAAAAGCACC 3’). The PCR products were then subjected to double size selection to remove genomic DNA and primer dimer using AmPure XP beads (Beckman Coulter, A63881). 5% of the product was run on an agarose gel to ensure proper product size. The PCR products were then library-prepped and sequenced by Genewiz/Azenta Life Sciences using their Amplicon-EZ platform. Libraries were subjected to 2×250 bp Illumina sequencing. MaGeCK was used to calculate the read count of each individual sgRNA with no mismatches and compared to the sequence of the reference sgRNA. The read counts from the PCR amplified DNA isolated from the *in vivo* tumors were normalized to the read counts of the parental cell lines. MaGeCK was then used to identify positively enriched sgRNAs in *KP-H3^K36M^-Ctrl* + sgRNA pool injected mice relative to *KP-H3^WT^-Ctrl* + sgRNA pool injected mice using the default settings.

### Human dataset analysis

For gene expression analysis in human lung adenocarcinoma, RNA-sequencing data were obtained from the Cancer Genome Atlas lung adenocarcinoma dataset. All genes were ranked in order according to the genes most positively correlated with *Nsd2* expression to most negatively correlated, and this rank list was used to perform Gene Set Enrichment Analysis (GSEA) examining GO biological processes using the molecular signature database (MSigDB). Kaplan–Meier survival curves for lung adenocarcinoma patients with tumors expressing low (*n* = 581) or high (*n* = 580) levels of *Nsd2* were obtained from KM plotter (kmplot.com)

### Statistics and Reproducibility

All statistical analyses were performed using GraphPad Prism 10. For violin plots, unpaired Student’s t-tests were performed. For tumor volume tracking two-way ANOVA was performed. Sample sizes for individual experiments are indicated in the figures. Reproducibility of the findings was confirmed by analyzing at least n=3 technical and/or biological replicates. The findings of all replicates were consistent.

## ACKNOWLEDGEMENTS

We would like to thank ULAR staff for animal husbandry, J.Palma, K. Bennett, and R. Ly at the Molecular Pathology and Imaging Core (MPIC) for histological analysis and technical expertise, K. Ge and B. Capell for gifting the *LSL-H3^K36M^* transgenic mice, K. Budinich for help with ChIP, and M. Winslow and C. Li for helpful discussions and critical reading of the manuscript. This work was supported by NIH grants (R01-CA262619, R01-CA279698, R01-CA276350 to DMF, 2-T32-CA-15299-15 to ACG) and Department of Defense grant (W81XWH2210742 and W81XWH2210008 to DMF).

## AUTHOR CONTRIBUTIONS

ACG, DMF, and KMA performed animal studies. ACG, KMA, MRR, and DAR performed histopathological analyses and quantification. CDP performed spatial analyses with supervision from SMS. SV performed next-generation sequencing analyses with supervision from IAA. ACG, AMCS, KMA, MRR, VIN, and NFF performed cell culture studies with supervision from DMF. S. Stransky performed histone mass spectrometry with supervision from S. Sidoli. DMF and ACG interpreted all datasets. DMF and ACG conceived and designed the study. DMF and ACG wrote the manuscript with editorial assistance from KMA, ACMS, and VIN.

---

**Extended Data Figure 1:**
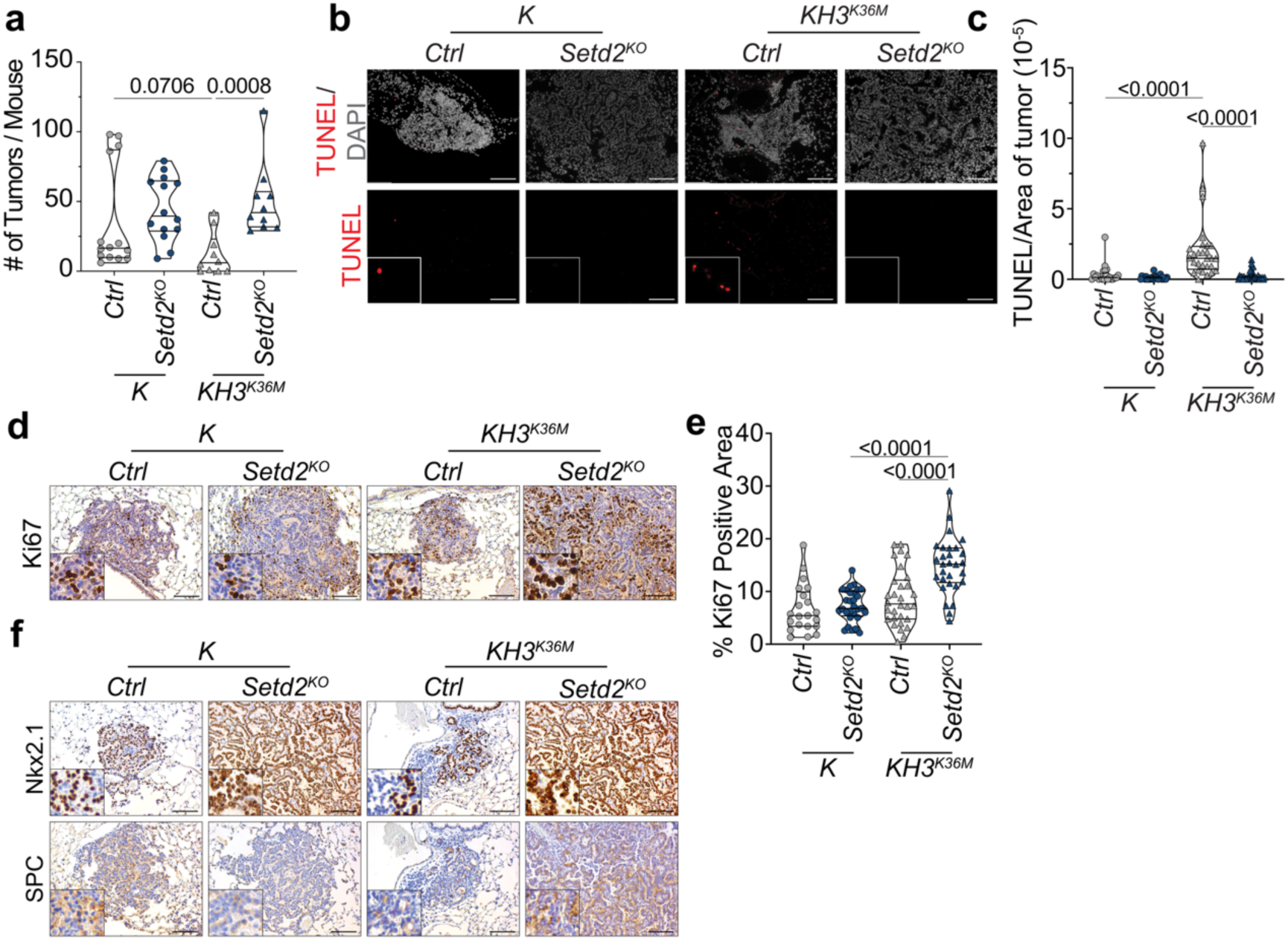
H3^K36M^ promotes tumor cell death in a Setd2-dependent manner. a. Quantification of tumor number in *K-Ctrl*, *K-Setd2^KO^*, *KH3^K36M^-Ctrl*, and *KH3^K36M^-Setd2^KO^* mice. Significance was determined by unpaired Student’s t-test. Error bars represent mean ±SEM. b. TUNEL staining of tumors from *K-Ctrl*, *K-Setd2^KO^*, *KH3^K36M^-Ctrl*, and *KH3^K36M^-Setd2^KO^* mice. Scale bars are 119 μm; insets are magnified x3 c. Quantification of TUNEL staining in *K-Ctrl*, *K-Setd2^KO^*, *KH3^K36M^-Ctrl*, and *KH3^K36M^-Setd2^KO^* tumors. Significance was determined by unpaired Student’s t-test. Error bars represent mean±SEM. d. IHC for Ki67 in *K-Ctrl*, *K-Setd2^KO^*, *KH3^K36M^-Ctrl*, and *KH3^K36M^-Setd2^KO^* tumors. Scale bars are 119 μm; insets are magnified x3. e. Quantification of Ki67 IHC staining in *K-Ctrl*, *K-Setd2^KO^*, *KH3^K36M^-Ctrl*, and *KH3^K36M^-Setd2^KO^* mice. Significance was determined by unpaired Student’s t-test. Error bars represent mean±SEM. f. IHC for lung lineage markers Nkx2.1 (TTF1) and SPC in *K-Ctrl*, *K-Setd2^KO^*, *KH3^K36M^-Ctrl*, and *KH3^K36M^-Setd2^KO^* tumors. Scale bars are 119 μm; insets are magnified x3.

**Extended Data Figure 2:**
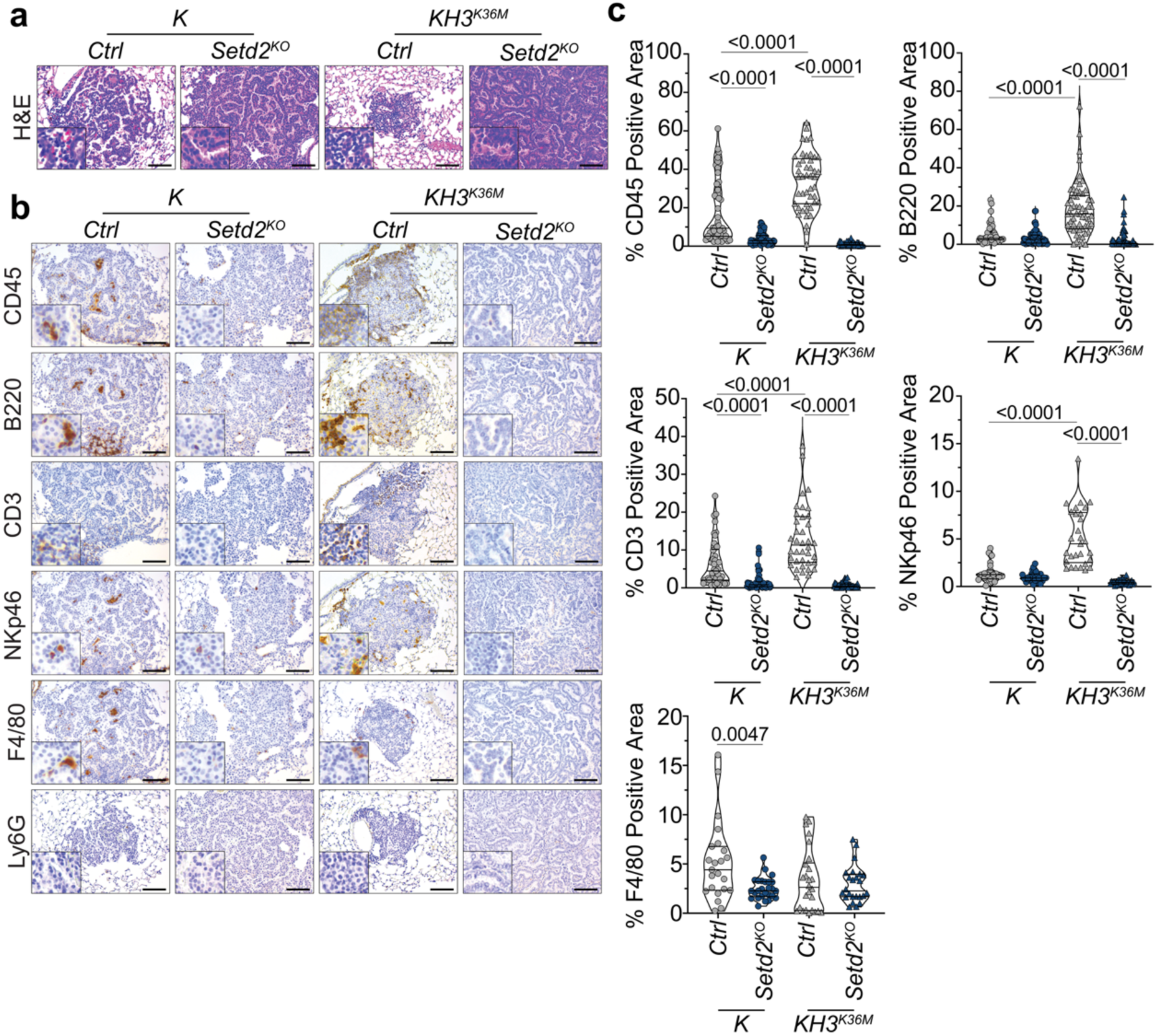
H3^K36M^ promotes Setd2-dependent immune infiltration. a. Representative H&E images of tumors from *K-Ctrl*, *K-Setd2^KO^*, *KH3^K36M^-Ctrl*, and *KH3^K36M^-Setd2^KO^* mice. Scale bars are 119 μm. b. IHC for immune cell markers CD45, B220, CD3, NKp46, F4/80, and Ly6G in *K-Ctrl*, *K-Setd2^KO^*, *KH3^K36M^-Ctrl*, and *KH3^K36M^-Setd2^KO^* tumors. Scale bars are 119 μm; insets are magnified x3. c. Quantification of CD45, B220, CD3, NKp46, and F4/80 IHC staining in *K-Ctrl*, *K-Setd2^KO^*, *KH3^K36M^-Ctrl*, and *KH3^K36M^-Setd2^KO^* mice. Significance was determined by unpaired Student’s t-test. Error bars represent mean±SEM.

**Extended Data Figure 3:**
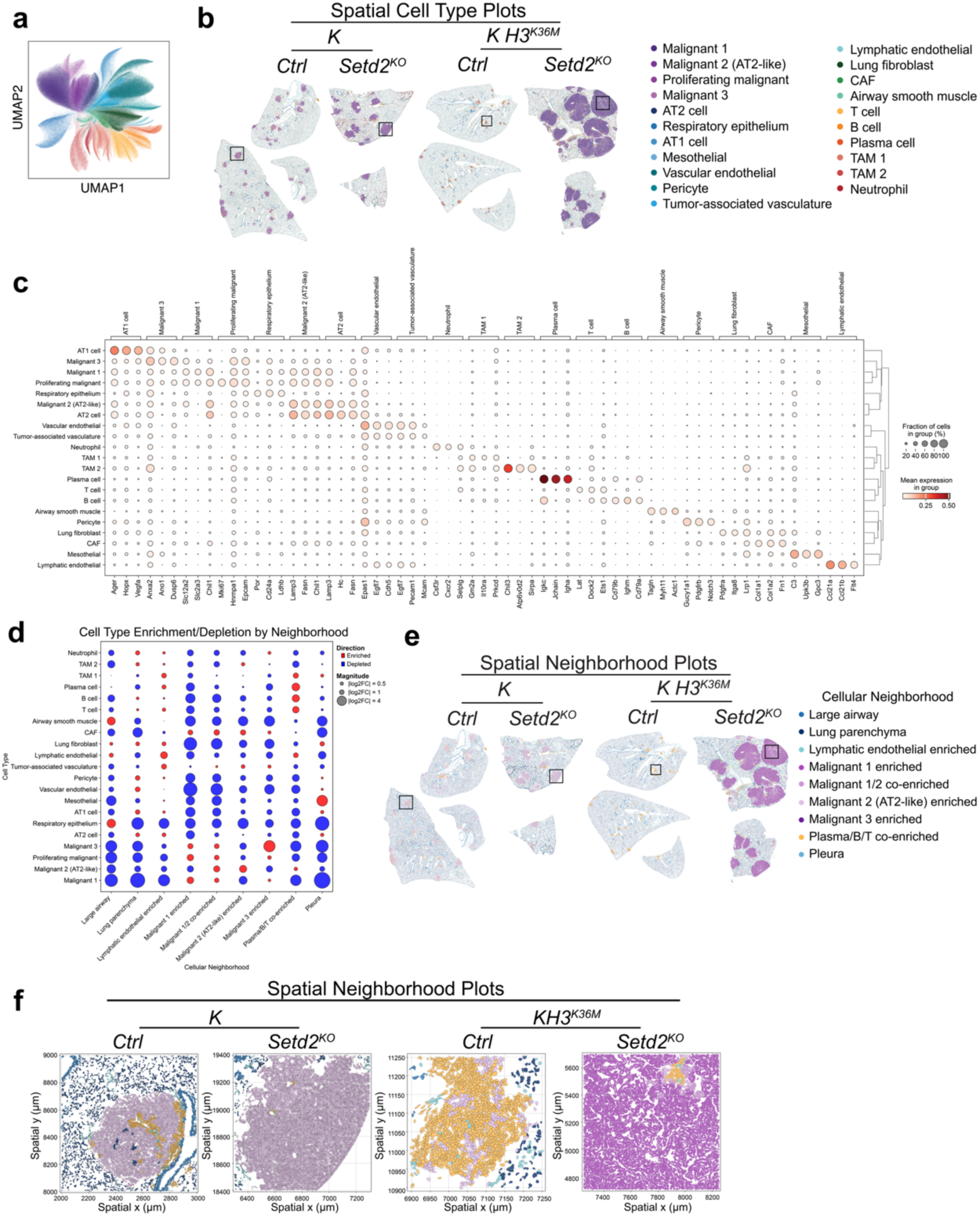
Spatial transcriptomics identifies H3^K36M^-promoted and Setd2-dependent immune infiltration and activation. a. Combined UMAP depicting cells from all four genotypes in **Fig. 2a**. b. Spatial cell type plots of full lung lobes used in spatial analysis, with boxes indicating representative tumors in **Fig 2a**. c. Marker gene expression in cell types identified in Xenium spatial analysis. d. Fold change enrichment or depletion of cell types by cellular neighborhood. e. Spatial cellular neighborhood plots of full lung lobes used in spatial analysis, with boxes indicating representative tumors in **Fig 2a**. f. Spatial cellular neighborhood plots of representative tumors from **Fig. 2a**.

**Extended Data Figure 4:**
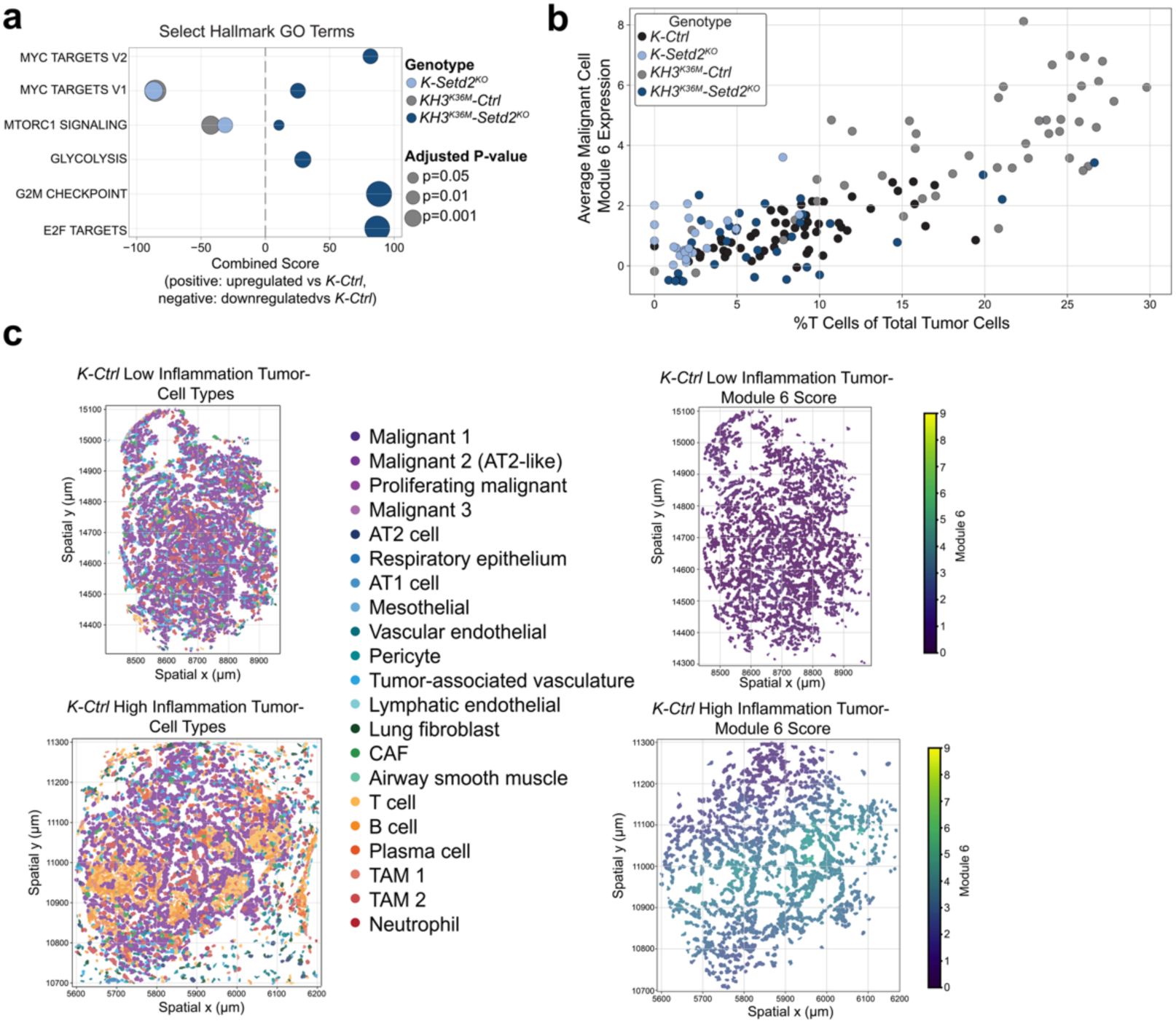
Module 6 expression correlates with inflammation. a. Gene Ontology enrichment scores for GO Terms associated with increased tumor aggressiveness. Enrichment scores were calculated for the sets of differentially upregulated and downregulated genes in *K-Setd2^KO^*, *KH3^K36M^-Ctrl*, and *KH3^K36M^-Setd2^KO^* tumors compared to *K-Ctrl*. Enrichment for a GO Term in differentially upregulated genes is depicted on the positive x-axis while enrichment in downregulated genes is depicted on the negative x-axis. Non-significant enrichments are not plotted. b. Scatter plot depicting T cell infiltration and mean Module 6 (inflammatory module) score in malignant cells on a per-tumor basis, colored by genotype. c. Cell types and Module 6 scores in T-cell low and T-cell high *K-Ctrl* tumors.

**Extended Data Figure 5:**
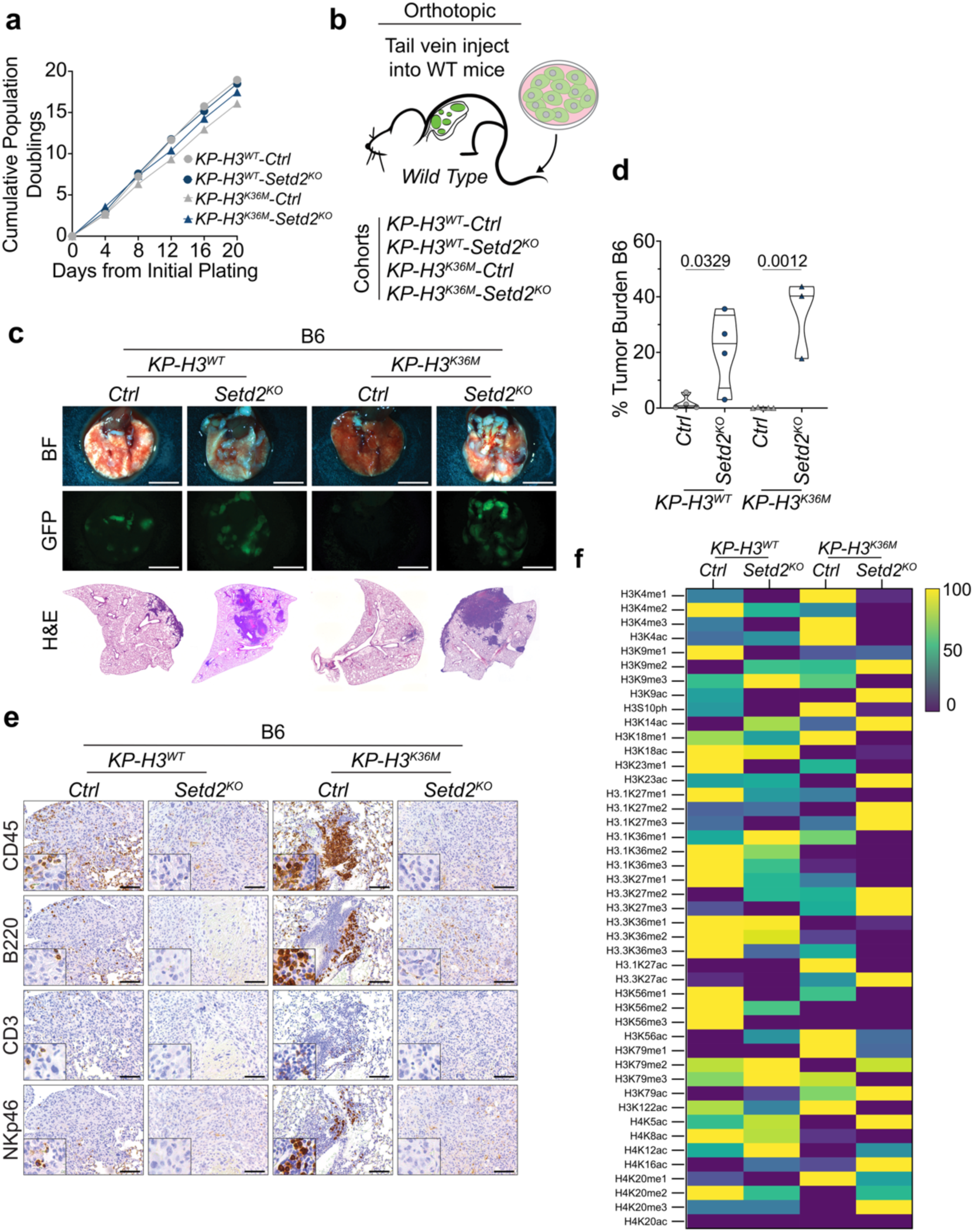
H3^K36M^-expressing cell lines phenocopy features of autochthonous tumors. a. Cell growth assay of *KP-H3^WT^-Ctrl*, *KP-H3^WT^-Setd2^KO^*, *KP-H3^K36M^-Ctrl*, and *KP-H3^K36M^-Setd2^KO^*cells. b. Experimental scheme of orthotopic tumor initiation. c. Brightfield and fluorescent micrographs of lungs from C57BL/J6 mice orthotopically injected with *KP-H3^WT^-Ctrl*, *KP-H3^WT^-Setd2^KO^*, *KP-H3^K36M^-Ctrl*, or *KP-H3^K36M^-Setd2^KO^*cells. Scale bars are 4.4 mm. Below are representative H&E images of tumor bearing lobes from C57BL/J6 mice orthotopically injected with *KP-H3^WT^-Ctrl*, *KP-H3^WT^-Setd2^KO^*, *KP-H3^K36M^-Ctrl*, or *KP-H3^K36M^-Setd2^KO^*cells. d. Quantification of tumor burden and individual tumor area in C57BL/J6 mice orthotopically injected with *KP-H3^WT^-Ctrl*, *KP-H3^WT^-Setd2^KO^*, *KP-H3^K36M^-Ctrl*, or *KP-H3^K36M^-Setd2^KO^* cells. Significance was determined by unpaired Student’s t-test. Error bars represent mean ±SEM. e. Representative images of IHC for immune cell markers CD45, B220, CD3, and NKp46 in tumors from C57BL/J6 mice orthotopically injected with *KP-H3^WT^-Ctrl*, *KP-H3^WT^-Setd2^KO^*, *KP-H3^K36M^-Ctrl*, or *KP-H3^K36M^-Setd2^KO^* cells. f. Relative abundance of histone modifications in *KP-H3^WT^-Ctrl*, *KP-H3^WT^-Setd2^KO^*, *KP-H3^K36M^-Ctrl*, and *KP-H3^K36M^-Setd2^KO^* cells by LC-MS/MS. Data are visualized in a row-normalized heat map.

**Extended Data Figure 6:**
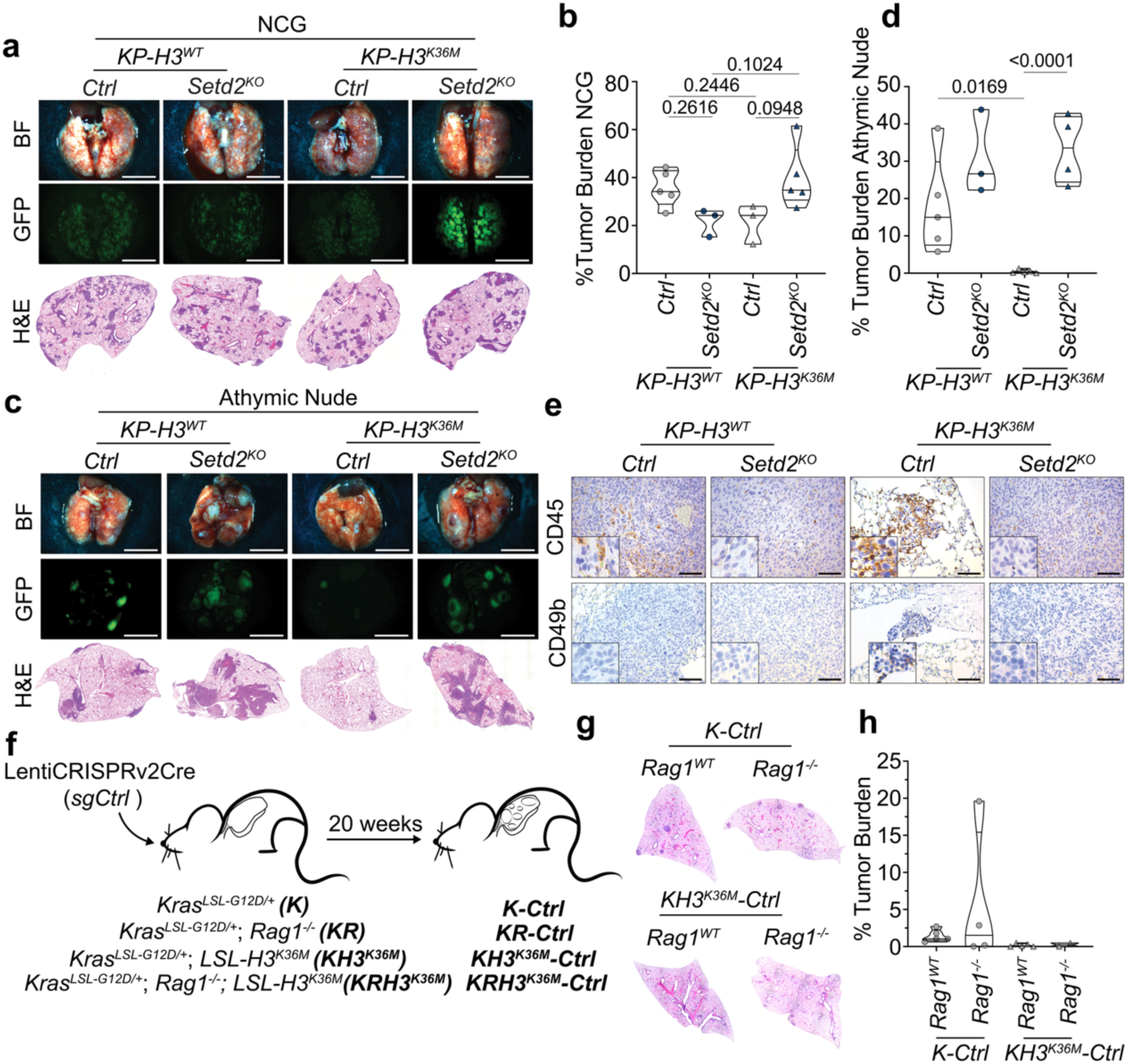
NK cells or T cells are sufficient for H3^K36M^-promoted cell death. a. Brightfield and fluorescent micrographs of lungs from NCG mice orthotopically injected with *KP-H3^WT^-Ctrl*, *KP-H3^WT^-Setd2^KO^*, *KP-H3^K36M^-Ctrl*, or *KP-H3^K36M^-Setd2^KO^*cells. Scale bars are 4.4 mm. Below are representative H&E images of tumor bearing lobes from NCG mice orthotopically injected with *KP-H3^WT^-Ctrl*, *H3^WT^-Setd2^KO^*, *H3^K36M^-Ctrl*, or *H3^K36M^-Setd2^KO^*cells. b. Quantification of tumor burden in NCG mice orthotopically injected with *KP-H3^WT^-Ctrl*, *KP-H3^WT^-Setd2^KO^*, *KP-H3^K36M^-Ctrl*, or *KP-H3^K36M^-Setd2^KO^* cells. Significance was determined by unpaired Student’s t-test. Error bars represent mean±SEM. c. Brightfield and fluorescent micrographs of lungs from athymic nude mice orthotopically injected with *KP-H3^WT^-Ctrl*, *KP-H3^WT^-Setd2^KO^*, *KP-H3^K36M^-Ctrl*, or *KP-H3^K36M^-Setd2^KO^*cells. Scale bars are 4.4 mm. Below are representative H&E images of tumor bearing lobes from athymic nude mice orthotopically injected with *KP-H3^WT^-Ctrl*, *KP-H3^WT^-Setd2^KO^*, *KP-H3^K36^M-Ctrl*, or *KP-H3^K36M^-Setd2^KO^* cells. d. Quantification of tumor burden in athymic nude mice orthotopically injected with *KP-H3^WT^-Ctrl*, *KP-H3^WT^-Setd2^KO^*, *KP-H3^K36^M-Ctrl*, or *KP-H3^K36M^-Setd2^KO^*cells. Significance was determined by unpaired Student’s t-test. Error bars represent mean±SEM. e. IHC for immune cell markers CD45 and CD49b in tumors from athymic nude mice orthotopically injected with *KP-H3^WT^-Ctrl*, *KP-H3^WT^-Setd2^KO^*, *KP-H3^K36M^-Ctrl*, or *KP-H3^K36M^-Setd2^KO^*cells. Scale bars are 119 μm; insets are magnified x3. f. Experimental scheme of autochthonous tumor initiation in *K, KRag1^-/-^, KH3^K36M^, and KRag1^-/-^H3^K36M^* mice. g. Representative images of tumor bearing lobes from *K-Ctrl, KRag1^-/-^-Ctrl, KH3^K36M^-Ctrl, and KRag1^-/-^H3^K36M^-Ctrl* mice. h. Quantification of tumor burden in *K-Ctrl, KRag1^-/-^-Ctrl, KH3^K36M^-Ctrl, and KRag1^-/-^H3^K36M^-Ctrl* mice. Significance was determined by unpaired Student’s t-test. Error bars represent mean±SEM.

**Extended Data Figure 7:**
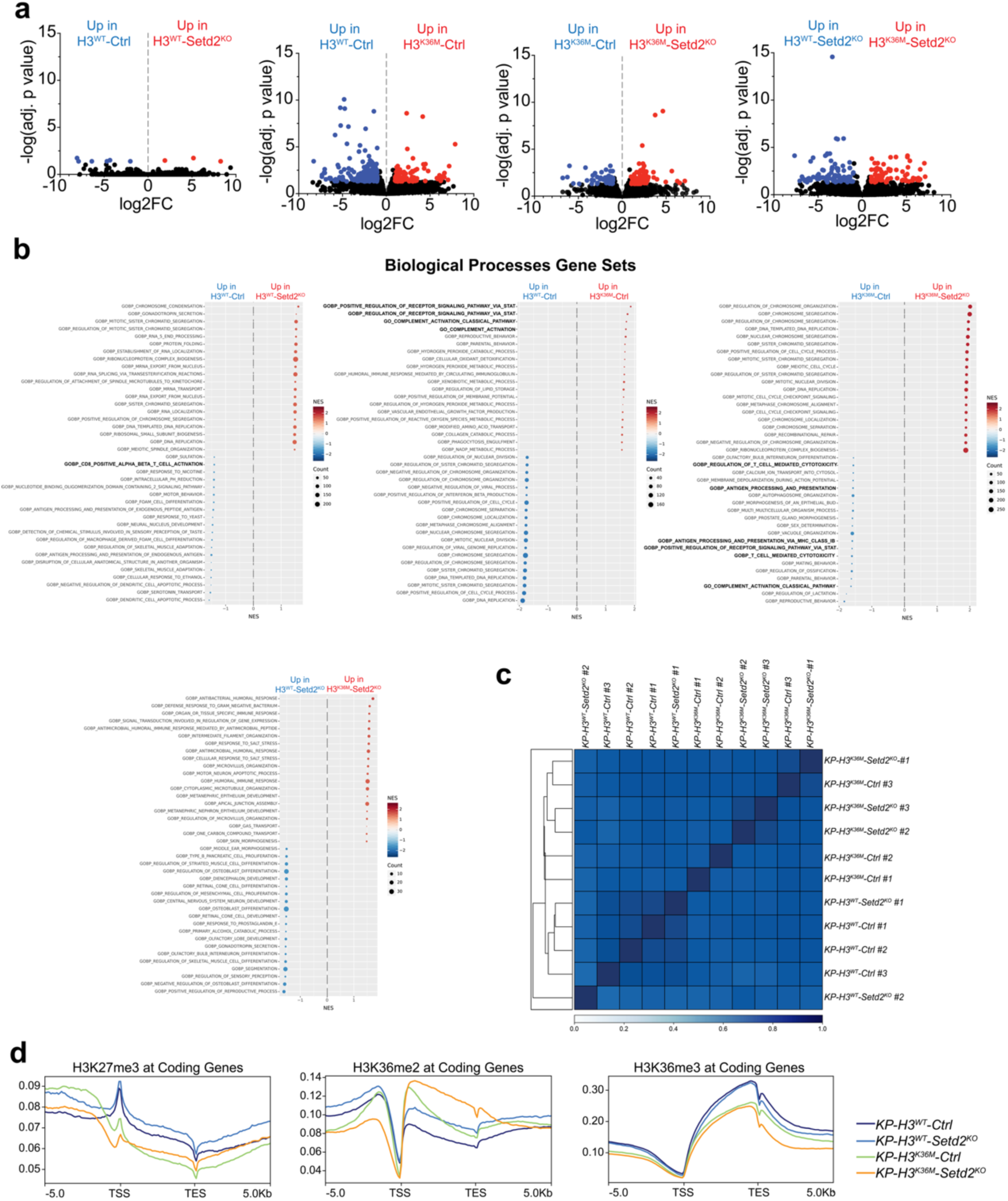
H3^K36M^ induces immune-related gene expression programs and remodels chromatin methylation. a. Volcano plots of RNA-sequencing data from *KP-H3^WT^-Ctrl*, *KP-H3^WT^-Setd2^KO^*, *KP-H3^K36M^-Ctrl*, and *KP-H3^K36M^-Setd2^KO^* cells. Comparisons are as listed. Colored dots represent genes that are differentially enriched (log2 fold-change greater-than 1 and false discovery rate (FDR)-adjusted P-value less-than 0.05) b. Bubble plot representation of top 20 up– and down-regulated GSEA biological processes gene sets in the noted comparisons. Normalized enrichment score (NES) plotted and the size of each dot represents the number of genes within the gene set. c. Correlation heatmap with hierarchical clustering amongst *KP-H3^WT^-Ctrl*, *KP-H3^WT^-Setd2^KO^*, *KP-H3^K36M^-Ctrl*, and *KP-H3^K36M^-Setd2^KO^*cell lines. d. Metagene plots describing the density of H3K27me3, H3K36me2, and H3K36me3 ChIP-seq across all coding genes. Plots are centered on coding genes elements across a ± 5 kb window.

**Extended Data Figure 8:**
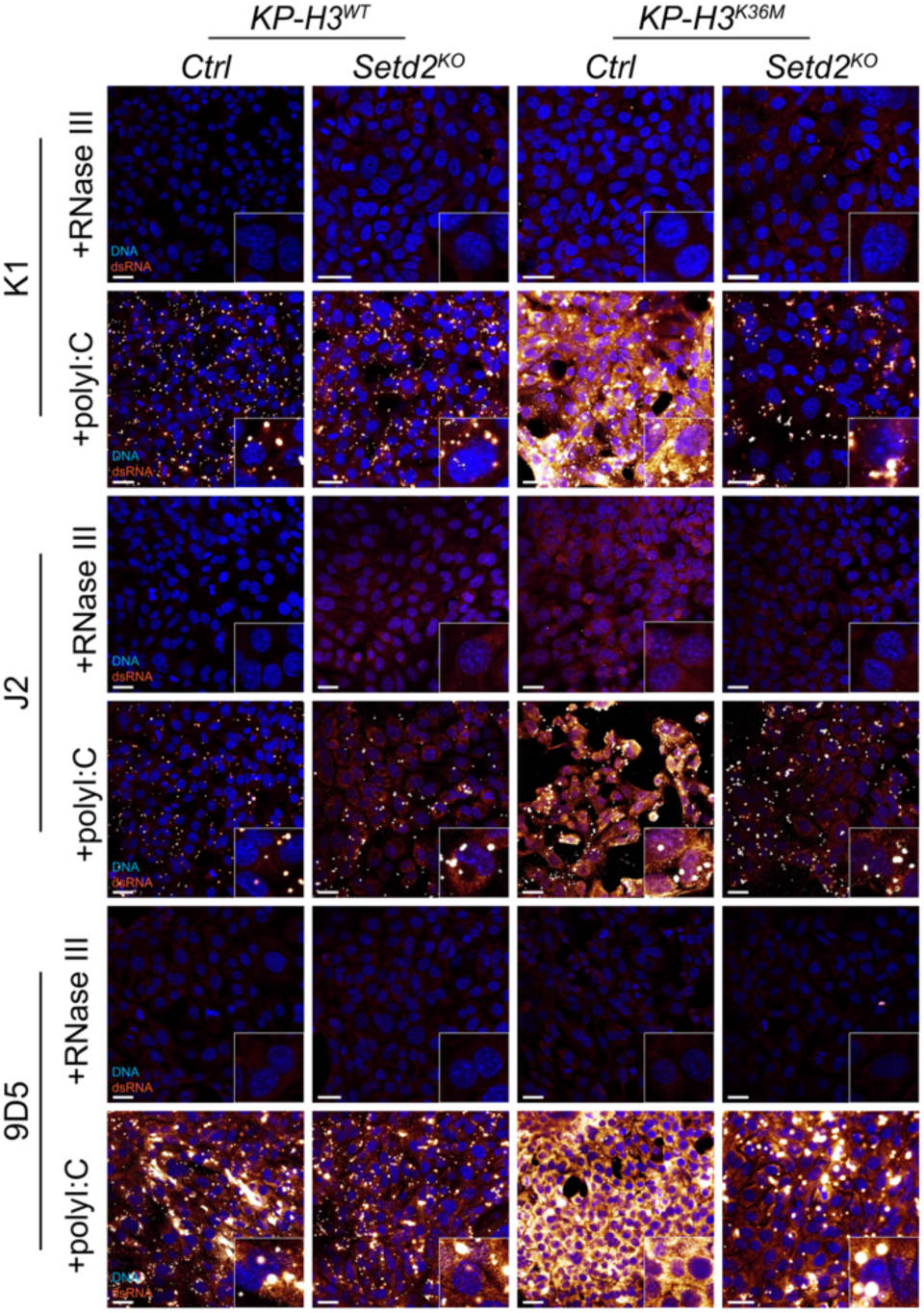
H3^K36M^-induced dsRNA signal is specific. a. Immunocytochemistry for dsRNA in *KP-H3^WT^-Ctrl*, *KP-H3^WT^-Setd2^KO^*, *KP-H3^K36M^-Ctrl*, and *KP-H3^K36M^-Setd2^KO^* cells. Staining was performed in parallel using 3 dsRNA antibodies: K1, J2, and 9D5. Cells received RNasee III treatment or were transfected with polyI:C.

**Extended Data Figure 9:**
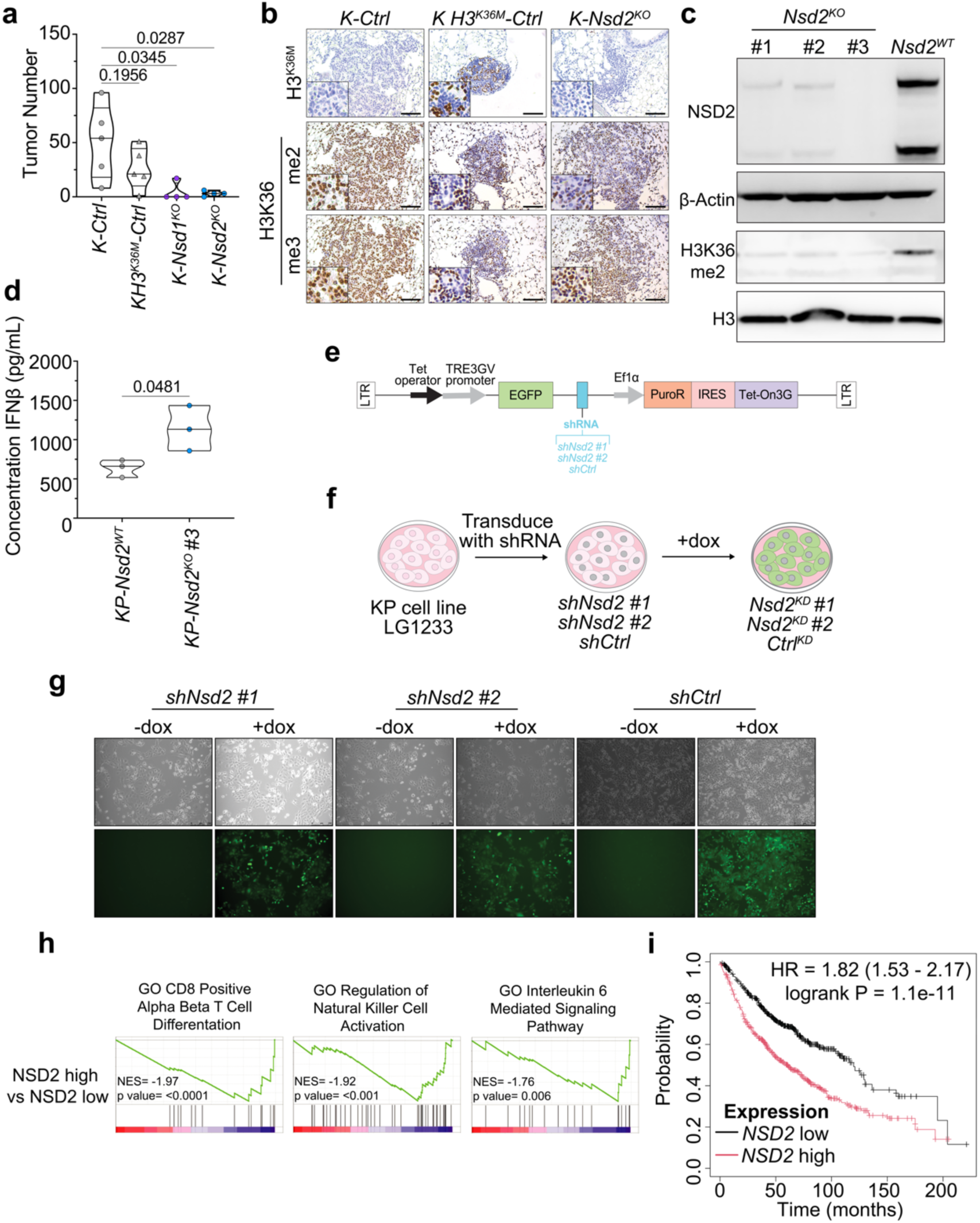
NSD2 promotes immune evasion in lung adenocarcinoma. a. Quantification of tumor number in *K-Ctrl*, *K-Setd2^KO^*, *KH3^K36M^-Ctrl*, and *KH3^K36M^-Setd2^KO^* mice. Significance was determined by unpaired Student’s t-test. Error bars represent mean ±SEM. b. IHC for H3^K36M^, H3K36me2, and H3K36me3 in *K-Ctrl*, *KH3^K36M^-Ctrl*, *K-Nsd1^KO^*, and *K-Nsd2^KO^* mice. Scale bars are 119 μm; insets are magnified x3. c. Immunoblot analysis of NSD2 and B-Actin in whole cell lysates and analysis of H3K36me2, H3K36me3, and H3 in histones prepared from *KP* and *KP-Nsd2^KO^* cells. d. IFN-β ELISA analysis of cell culture supernatant collected from *KP* parental and *KP-Nsd2^KO^* cells. Significance was determined by unpaired Student’s t-test. Error bars represent mean ±SEM. e. Schematic of plasmid used for doxycycline-induced shRNA expression. f. Experimental scheme for generation of *KP-*sh*Nsd2* and *KP-*sh*Ctrl* cell lines. g. Brightfield and fluorescent micrographs of doxycycline treated *KP-*sh*Nsd2* and *KP-*sh*Ctrl* cell lines. h. Select immune-related gene sets that are de-enriched in human lung adenocarcinoma patients with high *NSD2* expression. i. Kaplan–Meier survival curves for lung adenocarcinoma patients with tumors expressing low (*n* = 581) or high (*n* = 580) levels of *NSD2*.

**Extended Data Figure 10:**
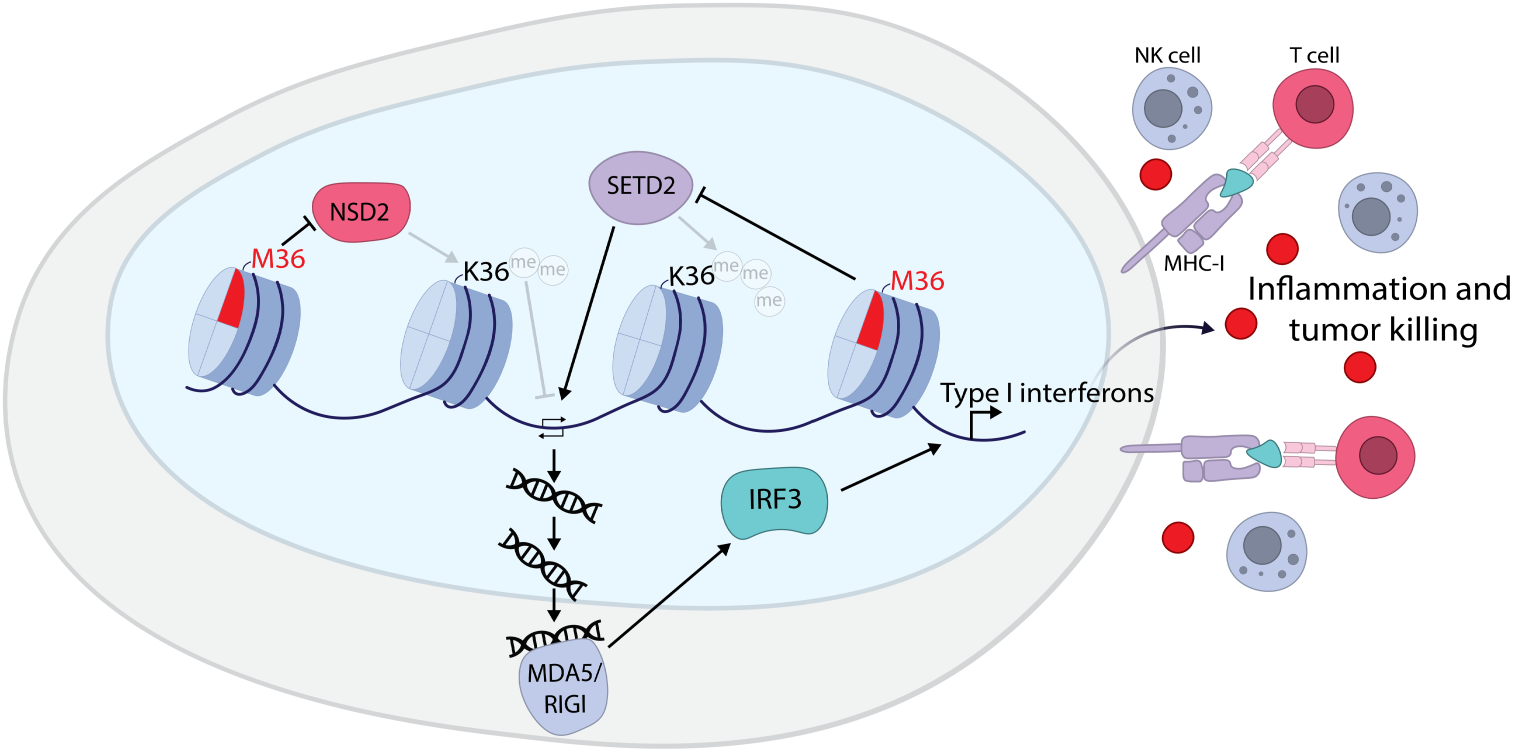
Loss of H3K36 dimethylation induces dsRNA expression that promotes Setd2-dependent, immune-mediated tumor killing. Expression of H3^K36M^ in tumor cells inhibits H3K36 methyltransferases SETD2 and NSD2. The resulting inhibition of H3K36me2 promotes dsRNA synthesis, which is detected by the MDA5/RIG-I pathway. Activation of this pathway promotes a strong inflammatory response, which leads to tumor cell killing. This entire process is dependent on intact SETD2, as inactivation of Setd2 prevents the formation of dsRNA species and the subsequent immune response is prevented.

**Supplementary Table 1:**
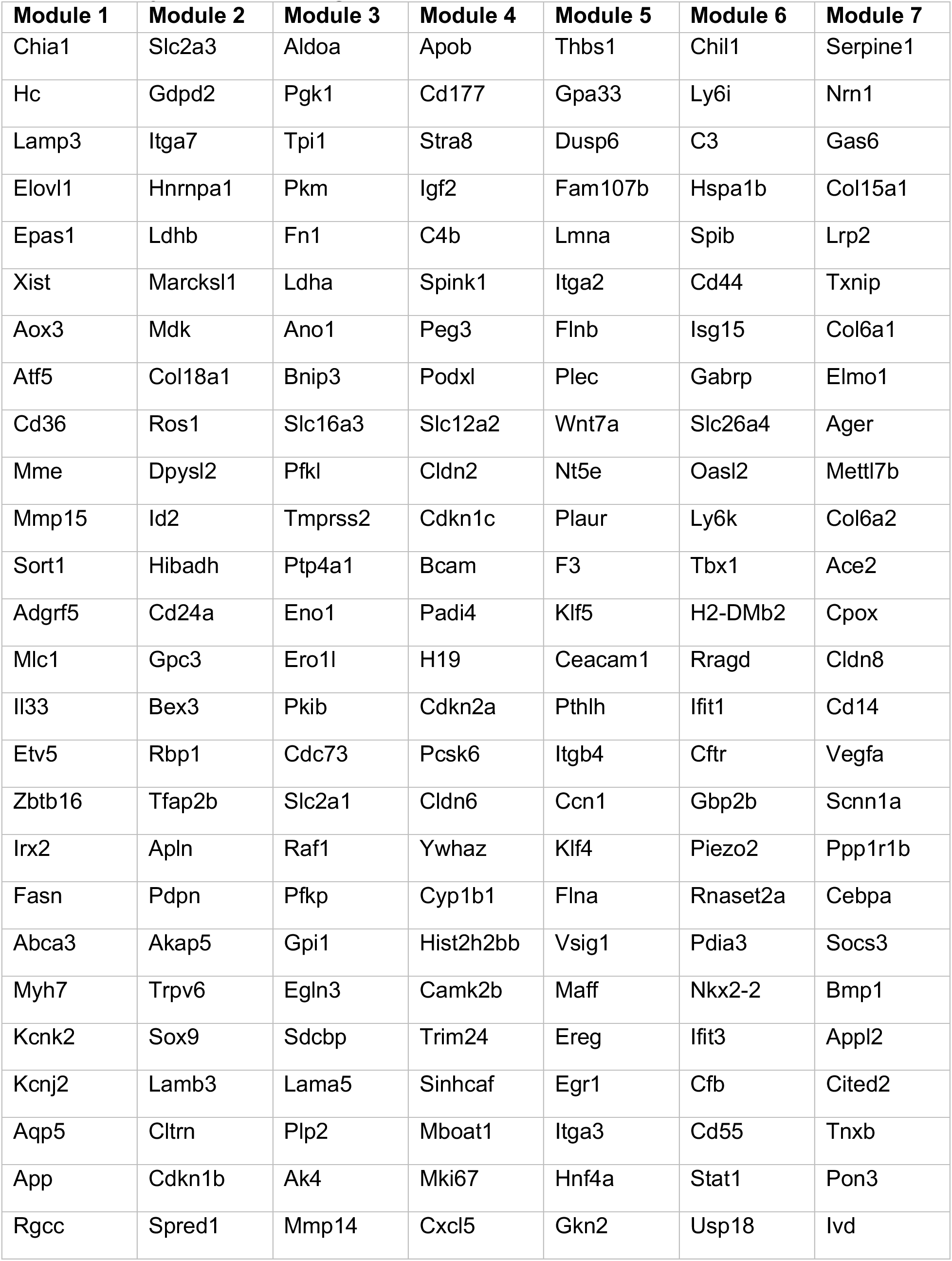
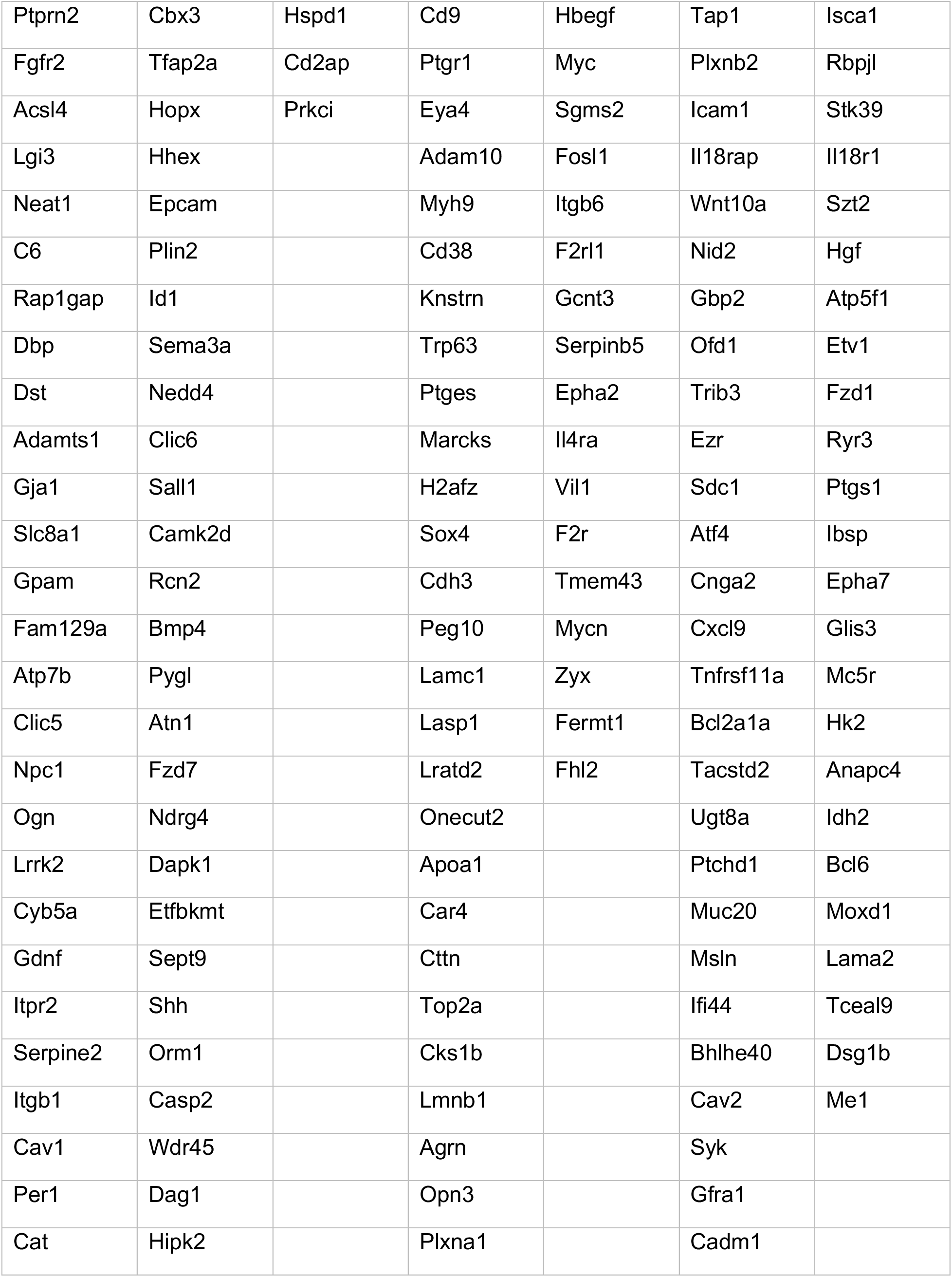
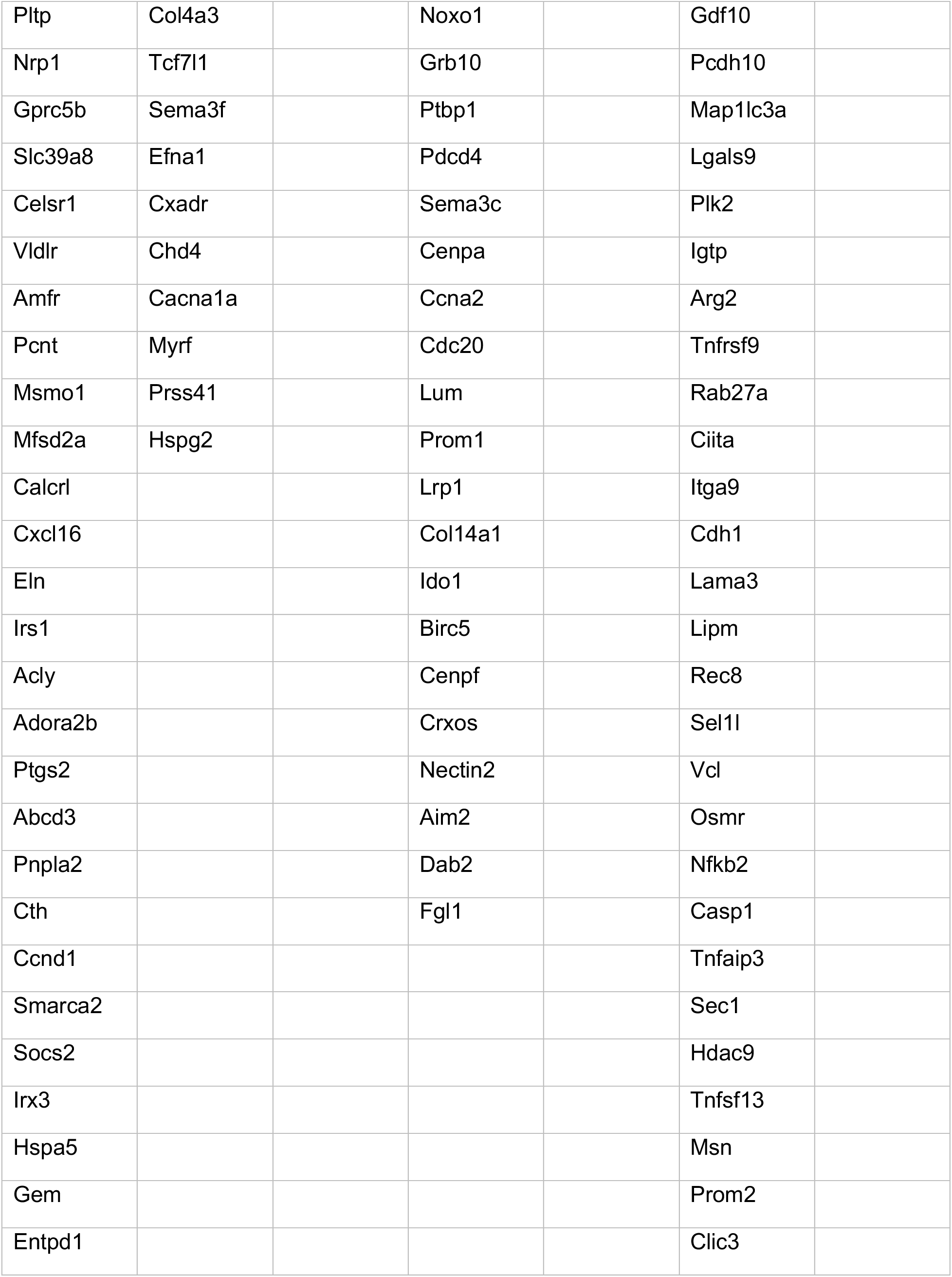
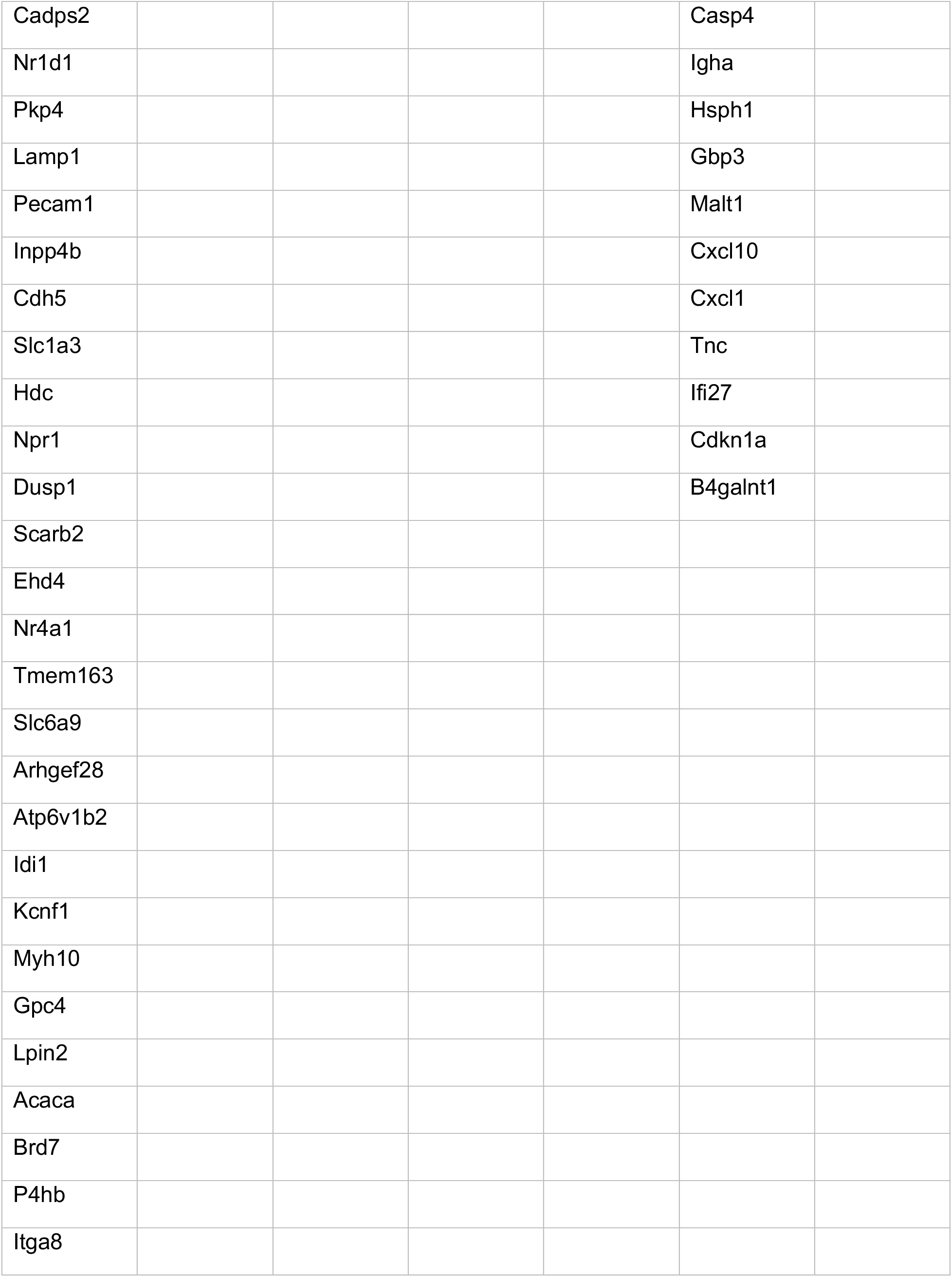
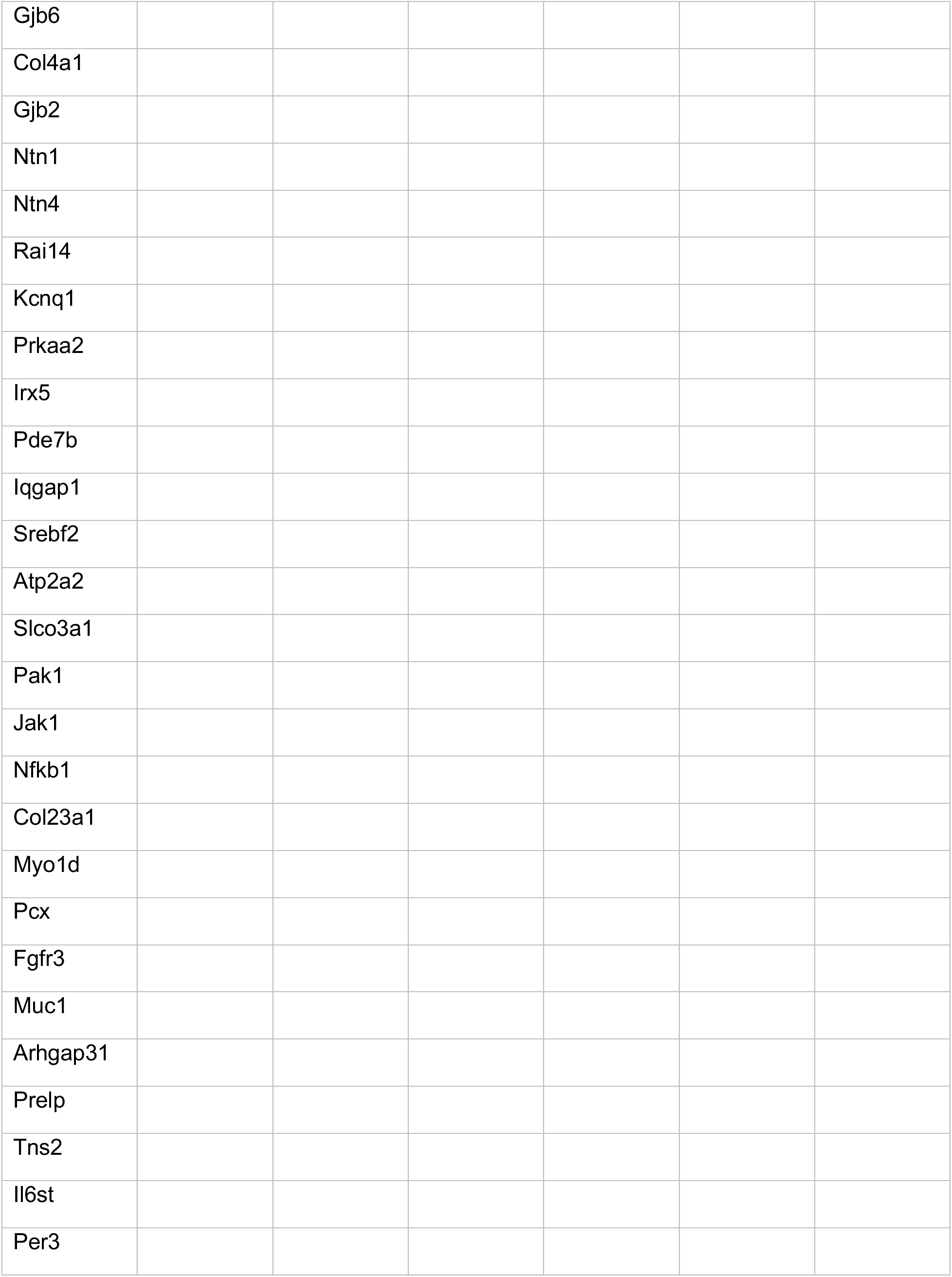
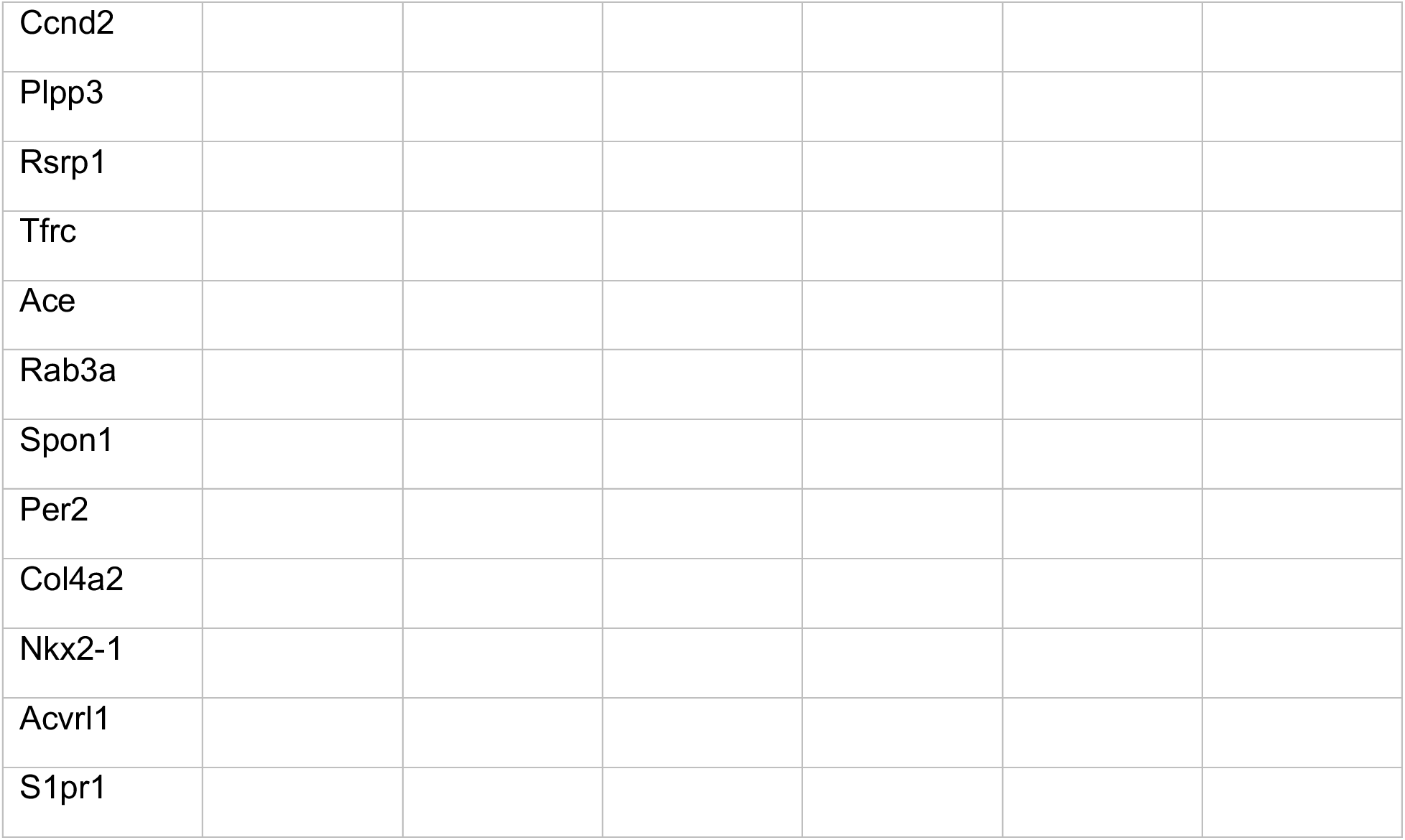
List of genes in Hotspot modules.

**Supplementary Table 2:**
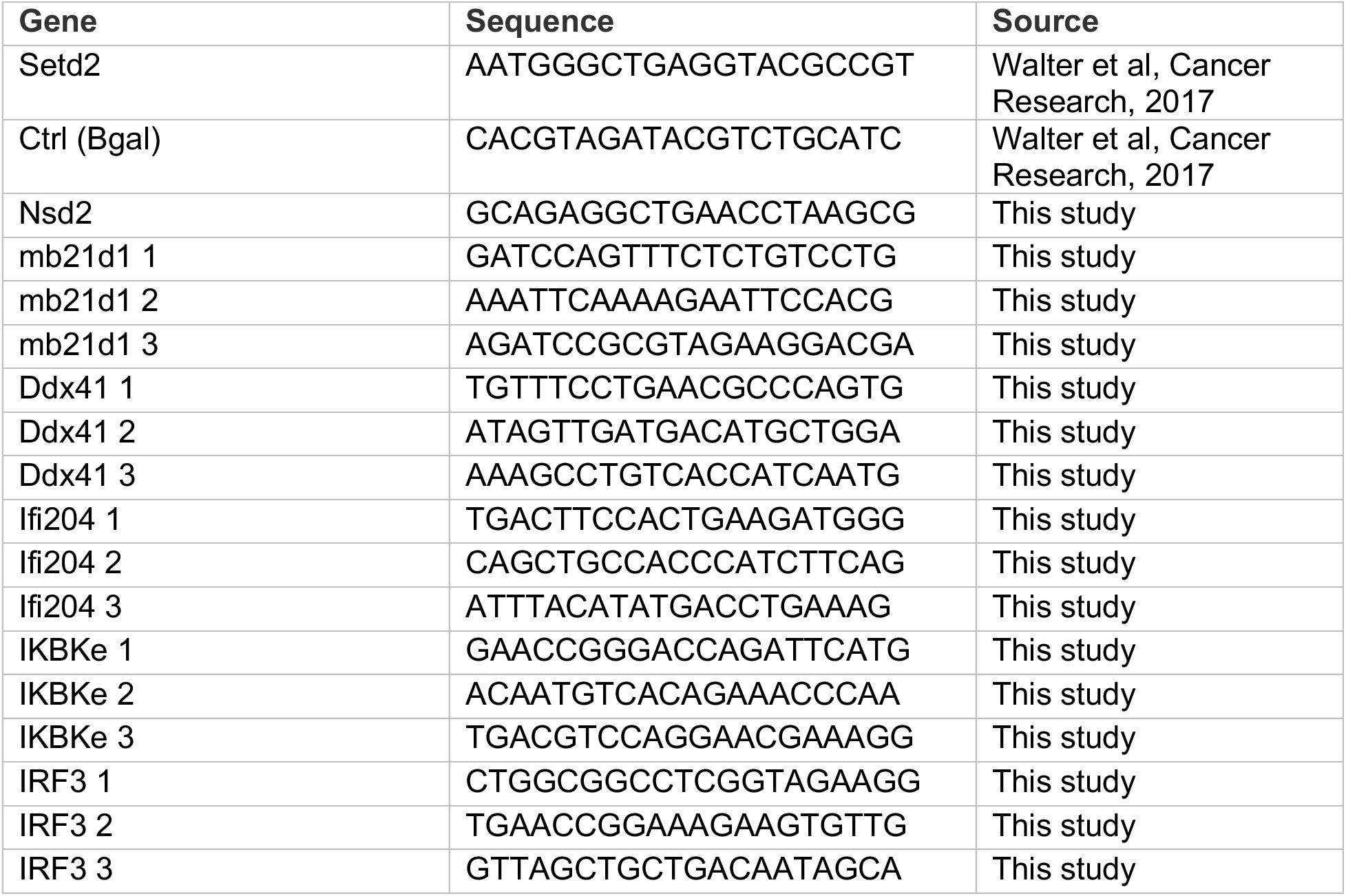
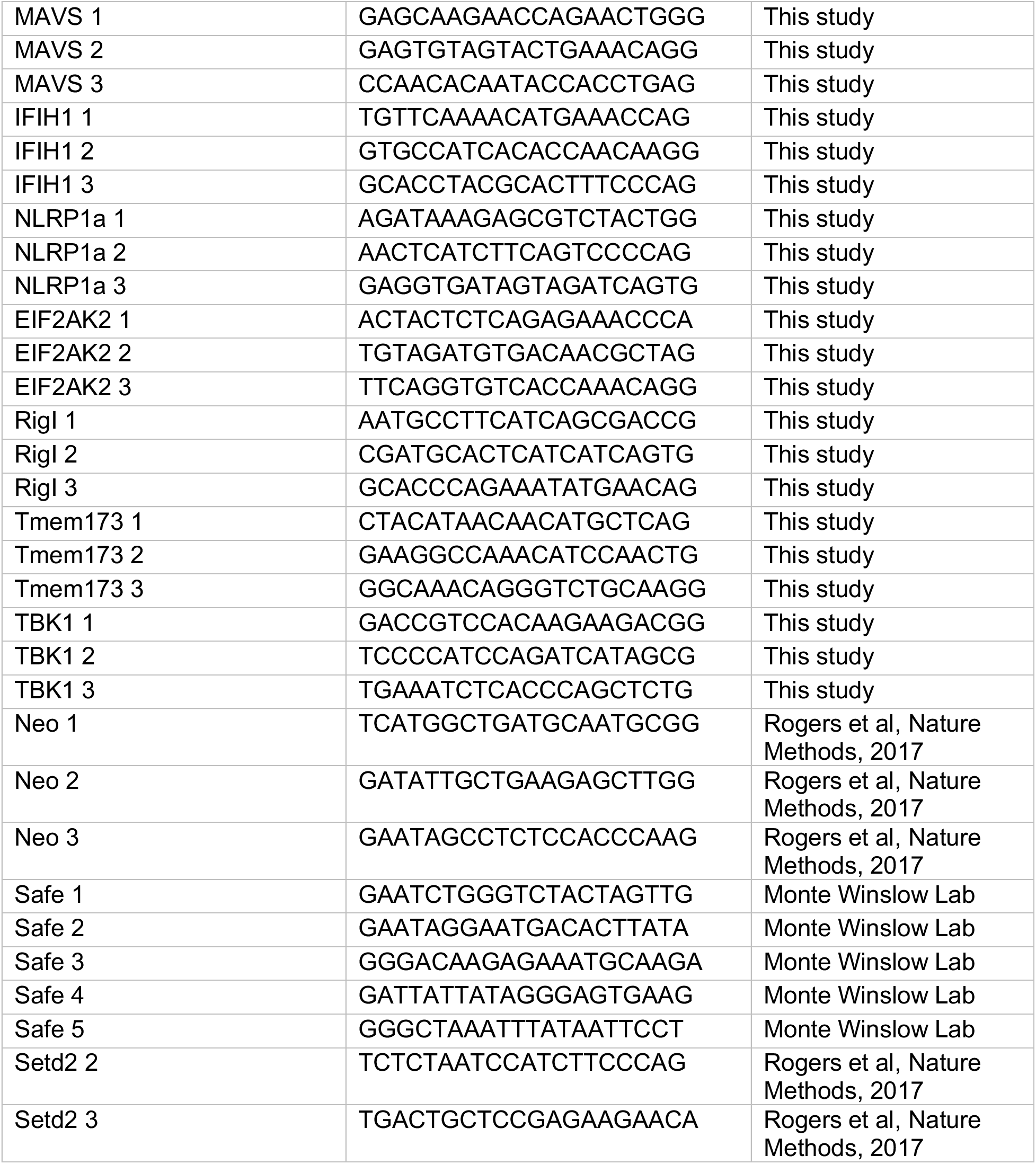
sgRNA sequences.

**Supplementary Table 3:**
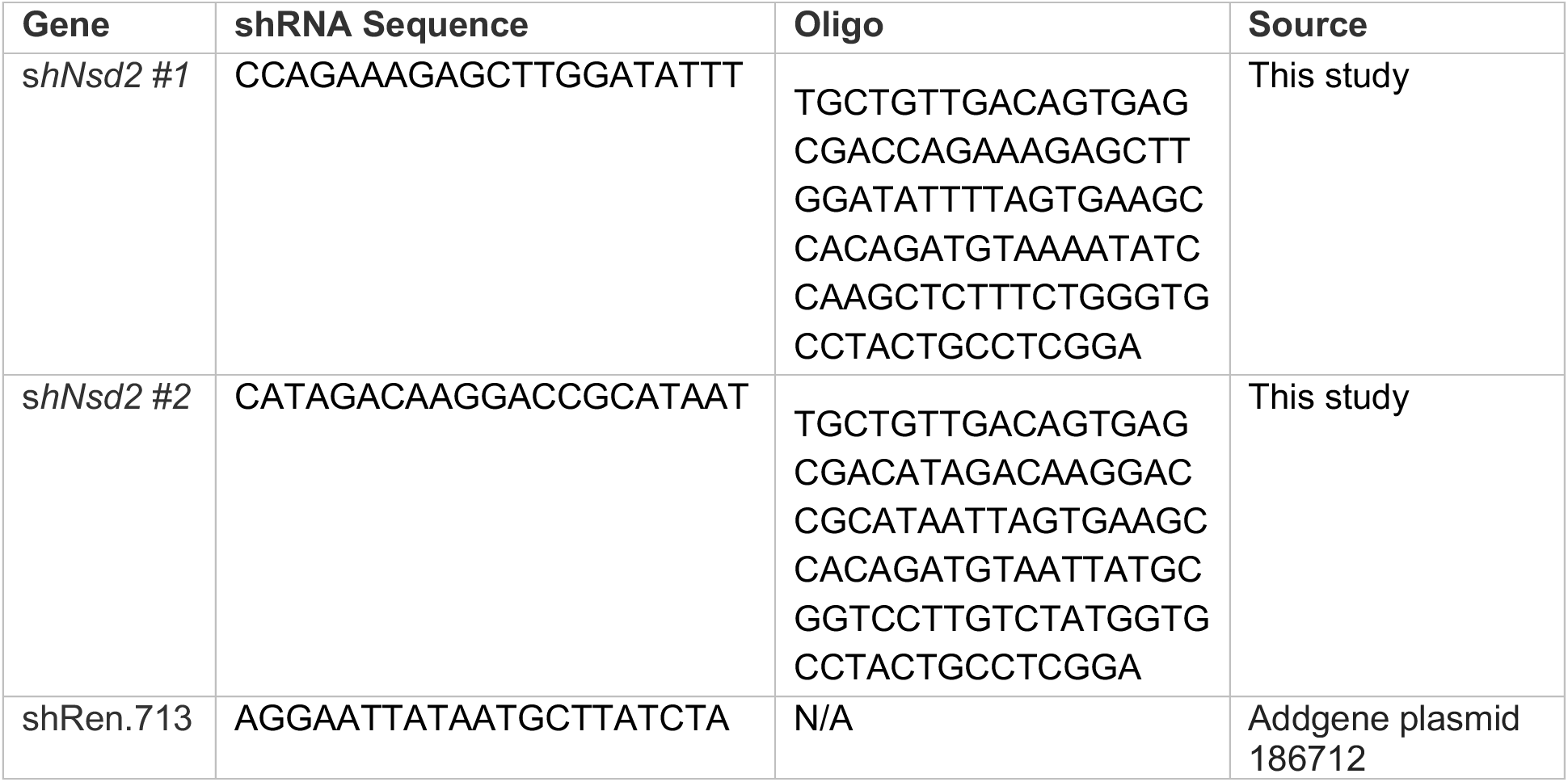
shRNA sequences.

**Supplementary Table 4:**
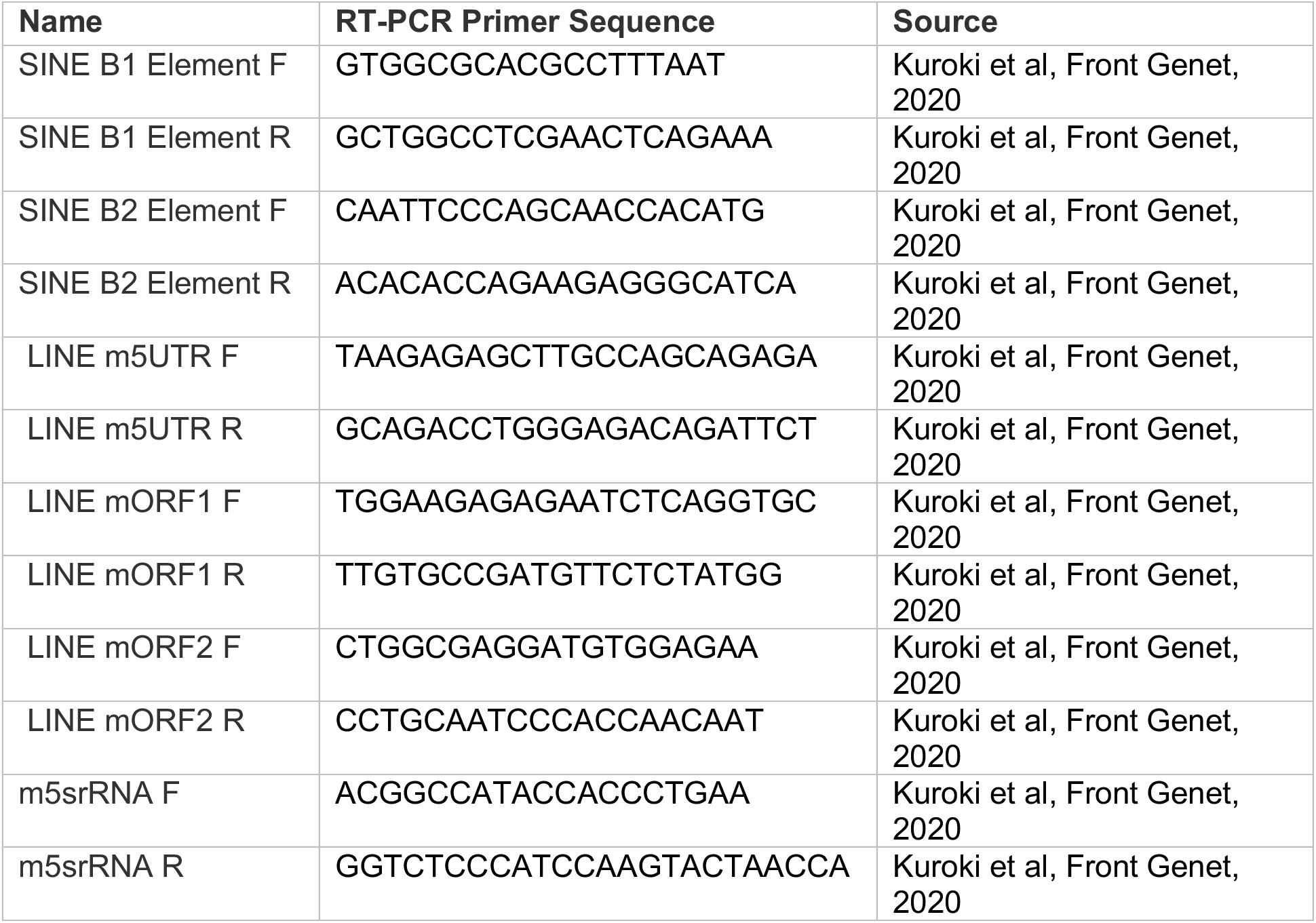
qRT-PCR primer sequences.

## REFERENCES

1. Sahu, V. and C. Lu, Oncohistones: Hijacking the histone code. Annu Rev Cancer Biol, 2022. 6: p. 293–312.

2. Fang, D., et al., The histone H3.3K36M mutation reprograms the epigenome of chondroblastomas. Science, 2016. 352(6291): p. 1344–8.

3. Zhang, Y., et al., Molecular basis for the role of oncogenic histone mutations in modulating H3K36 methylation. Sci Rep, 2017. 7: p. 43906.

4. Lam, U.T.F., et al., Structural and functional specificity of H3K36 methylation. Epigenetics Chromatin, 2022. 15(1): p. 17.

5. Hoetker, M.S., et al., H3K36 methylation maintains cell identity by regulating opposing lineage programmes. Nat Cell Biol, 2023. 25(8): p. 1121–1134.

6. Cerami, E., et al., The cBio cancer genomics portal: an open platform for exploring multidimensional cancer genomics data. Cancer Discov, 2012. 2(5): p. 401–4.

7. Rogers, Z.N., et al., A quantitative and multiplexed approach to uncover the fitness landscape of tumor suppression in vivo. Nat Methods, 2017. 14(7): p. 737–742.

8. Walter, D.M., et al., Systematic In Vivo Inactivation of Chromatin-Regulating Enzymes Identifies Setd2 as a Potent Tumor Suppressor in Lung Adenocarcinoma. Cancer Res, 2017. 77(7): p. 1719–1729.

9. Zehir, A., et al., Mutational landscape of metastatic cancer revealed from prospective clinical sequencing of 10,000 patients. Nat Med, 2017. 23(6): p. 703–713.

10. Walter, D.M., et al., Setd2 inactivation sensitizes lung adenocarcinoma to inhibitors of oxidative respiration and mTORC1 signaling. Commun Biol, 2023. 6(1): p. 255.

11. Kizer, K.O., et al., A novel domain in Set2 mediates RNA polymerase II interaction and couples histone H3 K36 methylation with transcript elongation. Mol Cell Biol, 2005. 25(8): p. 3305–16.

12. Rebehmed, J., et al., Expanding the SRI domain family: a common scaffold for binding the phosphorylated C-terminal domain of RNA polymerase II. FEBS Lett, 2014. 588(23): p. 4431–7.

13. Keogh, M.C., et al., Cotranscriptional set2 methylation of histone H3 lysine 36 recruits a repressive Rpd3 complex. Cell, 2005. 123(4): p. 593–605.

14. Dhayalan, A., et al., The Dnmt3a PWWP domain reads histone 3 lysine 36 trimethylation and guides DNA methylation. J Biol Chem, 2010. 285(34): p. 26114–20.

15. Naftelberg, S., et al., Regulation of alternative splicing through coupling with transcription and chromatin structure. Annu Rev Biochem, 2015. 84: p. 165–98.

16. Fahey, C.C. and I.J. Davis, SETting the Stage for Cancer Development: SETD2 and the Consequences of Lost Methylation. Cold Spring Harb Perspect Med, 2017. 7(5).

17. Huang, H., et al., Histone H3 trimethylation at lysine 36 guides m(6)A RNA modification co-transcriptionally. Nature, 2019. 567(7748): p. 414–419.

18. Bhattacharya, S., et al., The methyltransferase SETD2 couples transcription and splicing by engaging mRNA processing factors through its SHI domain. Nat Commun, 2021. 12(1): p. 1443.

19. Park, I.Y., et al., Dual Chromatin and Cytoskeletal Remodeling by SETD2. Cell, 2016. 166(4): p. 950–62.

20. Seervai, R.N.H., et al., The Huntingtin-interacting protein SETD2/HYPB is an actin lysine methyltransferase. Sci Adv, 2020. 6(40).

21. Chen, K., et al., Methyltransferase SETD2-Mediated Methylation of STAT1 Is Critical for Interferon Antiviral Activity. Cell, 2017. 170(3): p. 492–506 e14.

22. Yuan, H., et al., SETD2 Restricts Prostate Cancer Metastasis by Integrating EZH2 and AMPK Signaling Pathways. Cancer Cell, 2020. 38(3): p. 350–365 e7.

23. Yang, C., et al., Role of NSD1 as potential therapeutic target in tumor. Pharmacol Res, 2021. 173: p. 105888.

24. Sengupta, D., et al., NSD2 dimethylation at H3K36 promotes lung adenocarcinoma pathogenesis. Mol Cell, 2021. 81(21): p. 4481–4492 e9.

25. Fnu, S., et al., Methylation of histone H3 lysine 36 enhances DNA repair by nonhomologous end-joining. Proc Natl Acad Sci U S A, 2011. 108(2): p. 540–5.

26. Li, J., J.H. Ahn, and G.G. Wang, Understanding histone H3 lysine 36 methylation and its deregulation in disease. Cell Mol Life Sci, 2019. 76(15): p. 2899–2916.

27. Schmitges, F.W., et al., Histone methylation by PRC2 is inhibited by active chromatin marks. Mol Cell, 2011. 42(3): p. 330–41.

28. Janesick, A., et al., High resolution mapping of the tumor microenvironment using integrated single-cell, spatial and in situ analysis. Nat Commun, 2023. 14(1): p. 8353.

29. Schurch, C.M., et al., Coordinated Cellular Neighborhoods Orchestrate Antitumoral Immunity at the Colorectal Cancer Invasive Front. Cell, 2020. 182(5): p. 1341–1359 e19.

30. DeTomaso, D. and N. Yosef, Hotspot identifies informative gene modules across modalities of single-cell genomics. Cell Syst, 2021. 12(5): p. 446–456 e9.

31. Dimitrova, N., et al., Stromal Expression of miR-143/145 Promotes Neoangiogenesis in Lung Cancer Development. Cancer Discov, 2016. 6(2): p. 188–201.

32. Chen, Y.G. and S. Hur, Cellular origins of dsRNA, their recognition and consequences. Nat Rev Mol Cell Biol, 2022. 23(4): p. 286–301.

33. Rogers, Z.N., et al., Mapping the in vivo fitness landscape of lung adenocarcinoma tumor suppression in mice. Nat Genet, 2018. 50(4): p. 483–486.

34. Liu, Y., et al., Cryo-EM structure of SETD2/Set2 methyltransferase bound to a nucleosome containing oncohistone mutations. Cell Discov, 2021. 7(1): p. 32.

35. Jackson, E.L., et al., Analysis of lung tumor initiation and progression using conditional expression of oncogenic K-ras. Genes Dev, 2001. 15(24): p. 3243–8.

36. Srinivas, S., et al., Cre reporter strains produced by targeted insertion of EYFP and ECFP into the ROSA26 locus. BMC Dev Biol, 2001. 1: p. 4.

37. Zhuang, L., et al., Depletion of Nsd2-mediated histone H3K36 methylation impairs adipose tissue development and function. Nat Commun, 2018. 9(1): p. 1796.

38. DuPage, M., A.L. Dooley, and T. Jacks, Conditional mouse lung cancer models using adenoviral or lentiviral delivery of Cre recombinase. Nat Protoc, 2009. 4(7): p. 1064–72.

39. Gilbert, L.A., et al., Genome-Scale CRISPR-Mediated Control of Gene Repression and Activation. Cell, 2014. 159(3): p. 647–61.

40. Fang, Y., et al., The H3K36me2 methyltransferase NSD1 modulates H3K27ac at active enhancers to safeguard gene expression. Nucleic Acids Res, 2021. 49(11): p. 6281–6295.

41. Crowe, A.R. and W. Yue, Semi-quantitative Determination of Protein Expression using Immunohistochemistry Staining and Analysis: An Integrated Protocol. Bio Protoc, 2019. 9(24).

42. Sidoli, S., et al., Complete Workflow for Analysis of Histone Post-translational Modifications Using Bottom-up Mass Spectrometry: From Histone Extraction to Data Analysis. J Vis Exp, 2016(111).

43. Yuan, Z.F., et al., EpiProfile 2.0: A Computational Platform for Processing Epi-Proteomics Mass Spectrometry Data. J Proteome Res, 2018. 17(7): p. 2533–2541.

44. Patro, R., et al., Salmon provides fast and bias-aware quantification of transcript expression. Nat Methods, 2017. 14(4): p. 417–419.

45. Love, M.I., W. Huber, and S. Anders, Moderated estimation of fold change and dispersion for RNA-seq data with DESeq2. Genome Biol, 2014. 15(12): p. 550.

46. Subramanian, A., et al., Gene set enrichment analysis: a knowledge-based approach for interpreting genome-wide expression profiles. Proc Natl Acad Sci U S A, 2005. 102(43): p. 15545–50.

47. Kolde, R., pheatmap: Pretty Heatmaps. 2018.

48. Stephens, M., False discovery rates: a new deal. Biostatistics, 2016. 18(2): p. 275–294.

49. Wickham, H., ggplot2: Elegant Graphics for Data Analysis. 2016, Springer-Verlag New York.

50. Molder, F., et al., Sustainable data analysis with Snakemake. F1000Res, 2021. 10: p. 33.

51. Soneson, C., M.I. Love, and M.D. Robinson, Differential analyses for RNA-seq: transcript-level estimates improve gene-level inferences. F1000Res, 2015. 4: p. 1521.

52. Lawrence, M., R. Gentleman, and V. Carey, *rtracklayer: an R package for interfacing with genome browsers*. Bioinformatics, 2009. 25(14): p. 1841–2.

53. Langmead, B. and S.L. Salzberg, Fast gapped-read alignment with Bowtie 2. Nat Methods, 2012. 9(4): p. 357–9.

54. JM, G., Detecting sites of genomic enrichment. https://github.com/jsh58/Genrich, 2018.

55. Ramírez, F., et al., deepTools2: a next generation web server for deep-sequencing data analysis. Nucleic Acids Research, 2016. 44(W1): p. W160–W165.

56. Egan, B., et al., An Alternative Approach to ChIP-Seq Normalization Enables Detection of Genome-Wide Changes in Histone H3 Lysine 27 Trimethylation upon EZH2 Inhibition. PLoS One, 2016. 11(11): p. e0166438.

57. Jiacheng Chuan, A.Z., Lawrence Richard Hale, Miao He, Xiang Li, Atria, Atria: an ultra-fast and accurate trimmer for adapter and quality trimming. Gigabyte, 2021.

58. Perez, G., et al., The UCSC Genome Browser database: 2025 update. Nucleic Acids Res, 2025. 53(D1): p. D1243–D1249.

59. Ran, F.A., et al., Genome engineering using the CRISPR-Cas9 system. Nature protocols, 2013. 8(11): p. 2281–308.

